# Scattered differentiation of unlinked loci across the genome underlines ecological divergence of the selfing grass *Brachypodium stacei*

**DOI:** 10.1101/2023.06.06.543844

**Authors:** Wenjie Mu, Kexin Li, Yongzhi Yang, Adina Breiman, Jiao Yang, Ying Wu, Shuang Wu, Mingjia Zhu, Jianquan Liu, Eviatar Nevo, Pilar Catalan

## Abstract

Ecological divergence without geographic isolation, as an early speciation process that may lead finally to reproductive isolation through natural selection, remains one of the most interesting issues in evolutionary biology. However, the patterns of the underlying genetic divergences across the genome vary between different groups. Here we report that *Brachypodium stacei*, an inbreeding grass species, has been involved in sympatric ecological divergence without geographic isolation. Genomic, transcriptomic, and metabolomic analyses suggest that diploid *B. stacei* diverged sympatrically in two slopes with contrasting biomes at Evolution Canyon I (ECI), Mount Carmel, Israel, where gene flow has continued freely but reduced with the time. This ecological divergence involved the scattered divergence of many unlinked loci across the total genome that include both coding and non-coding regions. We also identified significantly differential expressions of ABA signaling pathway genes, and contrasting metabolome composition between the arid-*vs* forest-adapted *B. stacei* ECI populations. These results suggest that many small loci involved in environmental responses act additively to account for the ecological usages of this species in contrasted environments with gene flow.

**Significance:** Ecological divergence provides evidence for the origin of species through natural selection that has governed evolutionists’ attention since Darwin. In this study, we present multiple-omics analyses of two plant populations growing sympatrically in contrasted environments and revealed their distinct differentiation across all examined data. These two populations share the most recent ancestor compared with other populations and their divergence started in the early Holocene. We revealed that gene flow had continued but with a progressive reduction over time. The genetic divergences are scattered across the total genome involving many unlinked coding and non-coding regions. These findings highlight the significance of natural selection in the ecological divergence that may finally lead to species formation without geographic isolation.

## Introduction

The adaptation of ancestral populations to contrasted environments has been assumed to trigger ‘the origin of the species’ as a major mechanism since Darwin (1). Under divergent selection, two populations accumulate multiple phenotypic and physiological traits to adapt to differentiated niches even in the presence of gene flow (2). This divergence at the same site may occur across different species which further highlights the critical role of such an ecological selection in speciation (3). Although many empirical studies of the sympatric diverging populations based on genomic data were conducted, the genetic bases of niche usage remain largely inconsistent (3). The ecological divergence may involve genetic differentiation across the total genome with many unlinked additive loci because of the continuous gene flow (4). However, large genomic islands containing a number of linked genes with obvious selection signals have been reported in ecological divergences of a few species in a similar manner without geographic isolation (5). This is expected to be more distinct in inbreeding species because of the strong purifying selection and reduced diversity (6).

“Evolution Canyon I” (ECI), near Mount Carmel in Israel, is well known among evolutionary biologists as a hotspot of ecological divergence for diverse organisms, from bacteria to plants and mammals (7). Although the overall climate and geology are essentially identical, the microclimates of the two slopes of ECI strongly differ, with the south-facing ‘African’ slope (AS) receiving up to 300% higher solar radiation than the north-facing ‘European’ slope (ES) (Fig 1A) (8). Thus, the AS has a xeric biome that is tropical, hot, dry, and savannoid, whereas the ES is temperate, cool, humid, and forested. It is estimated that these distinctive microclimates have existed since the Plio-Pleistocene (5.3 Mya — 11.7 Kya) (8). Evidence of ecological divergence that may lead to sympatric speciation has been found for seven species from diverse organismic groups that are distributed on both slopes of ECI (7), including the haploid soil bacterium *Bacillus simplex* (9), diploid fruit-fly *Drosophila melanogaster* (10), grain beetle *Oryzaephilus surinamensis* (11), spiny mouse *Acomysca hirinus* (12-14), and wild barley *Hordeum spontaneum* (15), and polyploid crucifer *Ricotia lunaria* (16) and wild emmer wheat *Triticum dicoccoides* (17). The genomic divergence observed between AS and ES populations of these organisms has been found to correspond with allelic differentiation of genes that contribute to local adaptation to different environmental conditions.

**Figure 1.**
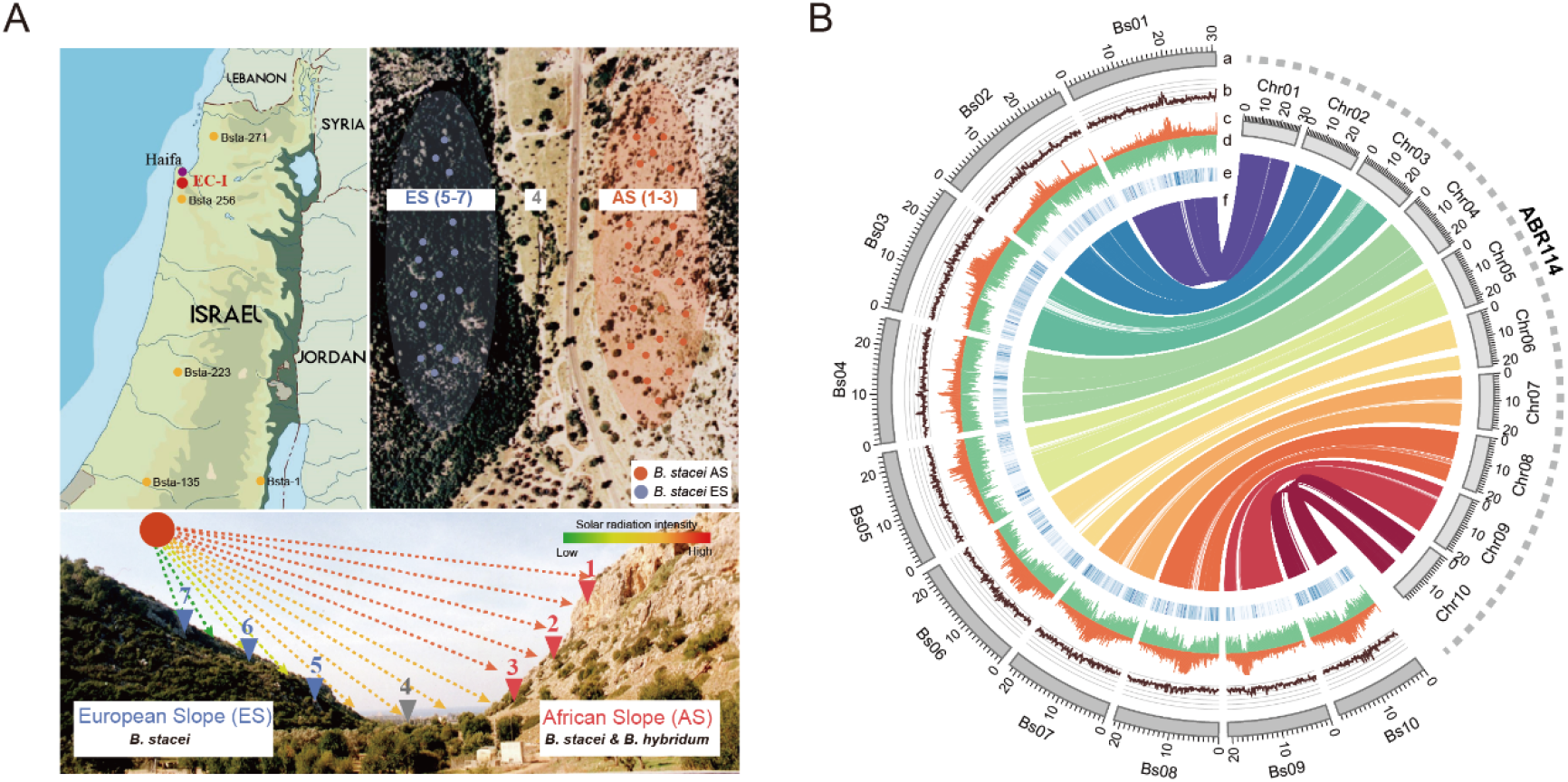
Sampling and genome organization of *Brachypodium stacei* ECI. **A)** Sampling locations of *B. stacei* in Evolution Canyon I (ECI) and other sites in Israel. Left-most panel: populations of *B. stacei* from ECI (red) and elsewhere in Israel (yellow). Right-most panel: populations of *B. stacei* ECI from the ‘European Slope’ (ES) (blue) and the ‘African Slope’ (AS) (salmon). Bottom panel: Despite being only 100-250 m apart in linear distance, these ES and AS slopes have strongly contrasted microclimates (e. g., solar radiation intensity differs substantially between them). **B)**. Overview of the newly assembled *B. stacei* genome from Evolution Canyon I (BstaECI, left) and syntenic relationships to the current *B. stacei* ABR114 v.1.1 reference genome (right). The tracks indicate: (a) chromosomes (Bs01-Bs10, BstaECI; JGI01-JGI10, ABR114), (b) GC contents, (c) transposable element densities, (d) gene models densities, (e) single nucleotide polymorphism (SNP) densities, and (f) collinearity of syntenic genes (BstaECI vs ABR114).

Here, we explore genetic bases underlying ecological divergence of an inbreeding grass species, *Brachypodium stacei* (2n=2x=20), which occurs on both the ES and AS in ECI (Fig. 1). This is an arid species that is broadly distributed in the southern part of the circum-Mediterranean region (18-20) where it grows in shady warm places or in open habitats, usually protected by shrubs (21-22). Using high-quality genome assemblies for *B. stacei*, genome-resequencing data from populations at ECI and surrounding regions in Israel, together with transcriptomic and metabolomic data, we tested whether genomic divergence occurred between the ES and AS populations of diploid *B. stacei*. We then examined how the highly divergent genes are functionally related and distributed across the total genome in this species in ECI, and how this model grass contributes to increase the organismal and biological extent of the sympatric speciation evolutionary scenario.

## Results

### Genomic assembly of *Brachypodium stacei* in ECI

The genome size of *B. stacei* ECI was estimated to be 263.59 Mb (with a low heterozygosity of 0.09%), which is ∼25Mb larger than the *B. stacei* reference genome ABR114 (Fig. S1). In order to make genomic analysis more reliable, an improved *de novo* assembly was generated for the *B. stacei* Bsta-ECI genome (Tables S1 and S2). The new *B. stacei* Bsta-ECI assembly had markedly higher contiguity than the current reference genome assembly *B. stacei* ABR114 v. 1.1 (https://phytozome-next.jgi.doe.gov/) (22), with ∼22.57Mb longer total contig length (256.71 Mb versus 234.17Mb), ∼98.5% lower total contig numbers (79 versus 3,132), and ∼44-fold higher N50 length (10.19 Mb vs 0.23 Mb) (Fig S2, Table S2). These high-quality contigs were further anchored onto 10 chromosomes assisted by 3D proximity information from Hi-C datasets (Table S3). The final genome assembly captured 248.99 Mb of the genome sequences, with 96.99% anchored percentages (Fig S3, Table S2). All chromosomes show longer sizes and fewer gaps than the reference genome ABR114 v. 1.1 (Table S4). The base call accuracy (QV) and assembly completeness were 39.78 and 99.18%, and Illumina pair-end reads mapping also showed a > 98.5% mapping rate and coverage rate than in the Bsta-ECI genome assembly (Table S5). 98.6% of the BUSCO ortholog and homeolog genes could be completely predicted in Bsta-ECI, which were slightly higher than those in the previous assembly (Fig. S4). In addition, the LAI (LTR Assembly Index) of Bsta-ECI was higher than values for the previously reported ABR114 assembly (Fig. S2C), indicating that Bsta-ECI has higher long-terminal repeat (LTR) retrotransposon completeness. All assessments suggested high consistency and completeness of the Bsta-ECI genome assembly which showed great improvement in contiguity and repetitive sequence completeness compared with the reference genome assembly ABR114.

Around 103.46 Mb (40.31%) of the total *B. stacei* Bsta-ECI genome sequence was annotated as repetitive element sequences (Table S6), a percentage higher than that of the reference genome ABR114 (Table S7). Most of the repetitive element sequence is Transposable elements (TEs), which include SINE (∼0.59%), LINE (∼7.33%), LTR (∼44.92%) and DNA (∼17.91%)

(Table S7). We further predicted 32,951 high-confidence protein coding genes in the *B. stacei* Bsta-ECI genome, slightly higher than those previously reported in ABR114 v. 1.1 (Fig. S4, Table S8). The gene structure feature (average CDS length, exon length, exon number, and intro length) are similar to those of the *B. stacei* ABR114 v. 1.1 and *B. distachyon* Bd21 reference genome (Fig. S5, Table S8). More than 97% of the predicted genes of the *B. stacei* Bsta-ECI genome had homologs in public functional databases (Table S9). We also found that 99.1% BUSCOs could be completely detected (Fig S4), indicating the high completeness of the gene model annotation. Furthermore, 1,838 transcription factors encoding genes, belonging to 67 gene families representing 5.5% of the total predicted genes, were found in the *B. stacei* Bsta-ECI genome (Table S10). The high-quality chromosome-level assembly and annotation of our *B. stacei* Bsta-ECI genome (Fig 1B) provide robust foundations for investigating the evolutionary processes that gave rise to potential sympatric speciation events involving *B*.*stacei* in Evolution Canyon I.

### Genetic divergence of two *B. stacei* populations in ECI

To explore the potential adaptive evolution of *B. stacei* in ECI, we subjected 46 *B. stacei* individuals (41 from ECI, 5 from other regions of Israel) to whole-genome resequencing (Fig. 1A, 2A, DateSet S1). A total of 328.87 GB of clean Illumina data were obtained and mapped to the Bsta-ECI local reference genome, with an average of ∼27× coverage for each individual. More than 98.5% average mapping rate and average genome coverage were obtained for both AS and ES individuals, indicating high alignment accuracy (DataSet S2). After variant calling and quality filtering, 220,321 short INDELs and 722,351 high-quality SNPs were identified for downstream analysis (Table S11). The average INDEL and SNP densities were 0.89/kb and 2.91/kb, respectively. We first constructed a maximum-likelihood (ML) phylogenetic tree (rooted with *B. distachyon* Bd30) to evaluate the relationships among *B. stacei* lineages. The phylogenetic tree separated the ECI individuals into two strongly supported clades (designated AS and ES), corresponding to their spatial distribution at ECI (Fig. 1A, 2A). The *B. stacei* lineages from other localities of Israel were resolved as either sister to the ECI clade or more distantly related, followed by the more divergent western and central Mediterranean lineages. Strong support was obtained for all main split nodes, and phylogenetic divergence tended to reflect overall geographic distances (Fig. 2A). The robust sister relationship between the two ECI populations indicates that the split was likely primary, rather than a result of secondary contact after allopatric divergence between geographically isolated populations (Figs. 1A, 2A). Principal component analysis (PCA) also showed that individuals collected from the ES slope clustered together and were separated from AS individuals along the first principal component axis (Fig. S6A). Population structure analysis for the best K=2 hypothetical populations was also consistent with the ML tree and PCA results (Fig. 2A, Fig. S6B). Interestingly, we observed a few genomically admixed individuals in both AS and ES populations, suggesting that some gene flow may still occur between these spatially adjacent populations (Fig. 2A, Fig. S6B).

**Figure 2.**
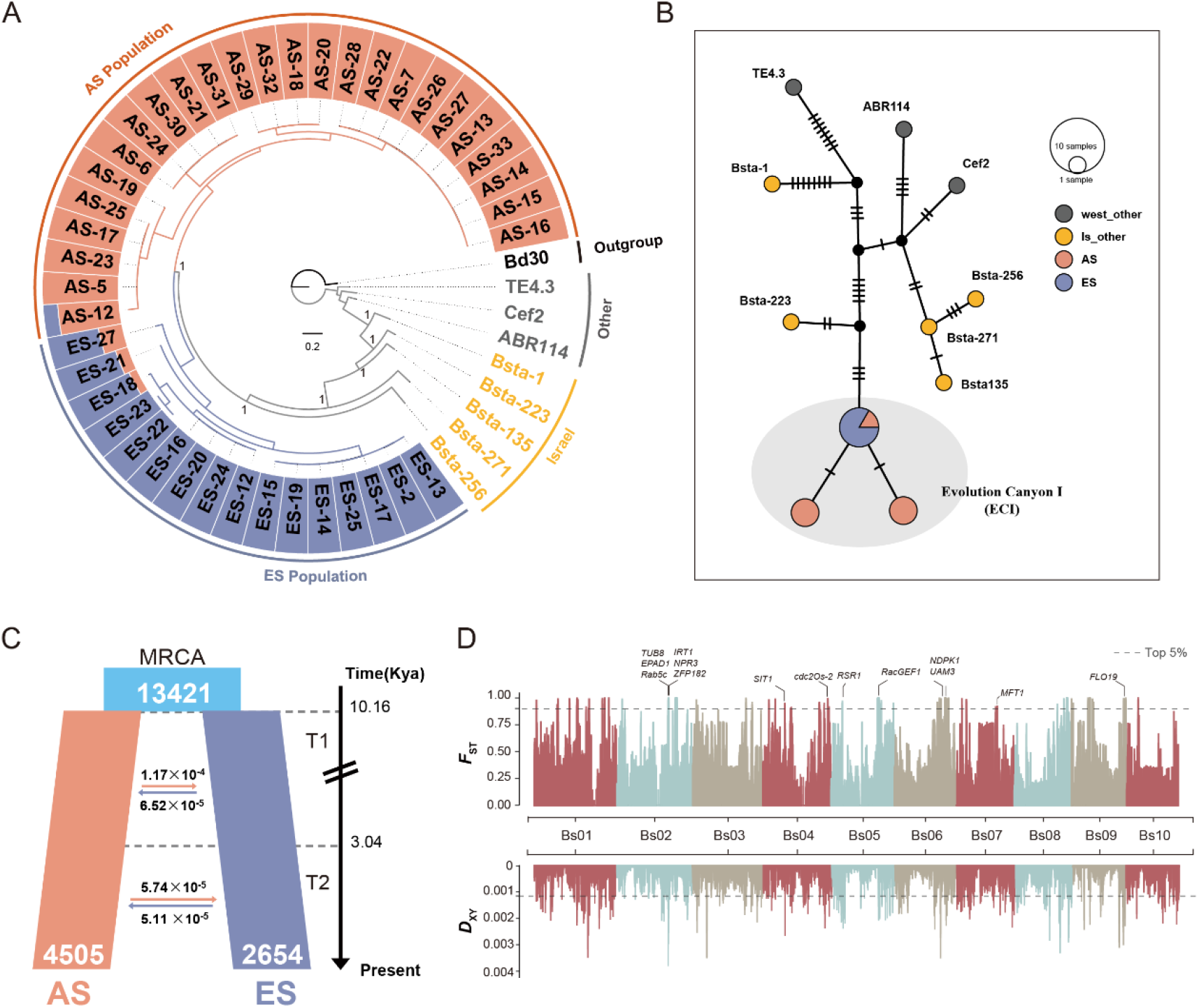
Genome divergence of *Brachypodium stacei* resequenced accessions from Evolution Canyon (ECI). **A)** Maximum-likelihood phylogenetic tree of the *B. stacei* accessions based on high-quality SNP data from individual genomes of *B. stacei* [AS (salmon), ES (blue), Israel (yellow), and other western and central Mediterranean populations (gray)], rooted with *B. distachyon* Bd30. The tree’s outer ring displays the population structure (with optimal K=2) of the *B. stacei* AS and ES populations. **B)** Statistical parsimony plastome haplotype network of *B. stacei* samples from Israel and other Mediterranean localities. The area of each circle in the network is proportional to the haplotype frequency, and the number of mutational steps between two nodes is indicated by short bars. **C)** Demographic history of the *B. stacei* AS and ES populations inferred by the best fit *∂a∂i* model (“asym_mig_twoepoch” model), indicating that a single split of the ancestral population (MRCA; light blue) 10.16 Ka gave rise to the modern AS and ES populations. The reciprocal average migration rates between ES and AS populations in two temporal epochs (T1 and T2) are shown by horizontal arrows and the estimated ages of T1 and T2 on the right side of the panel. **D)** Genetic differentiation and divergence between the *B. stacei* AS and ES populations revealed by *F*_*ST*_ and *D*_*XY*_ data. Dashed horizontal lines depict the top 5% thresholds (*F*_*ST*_ 0.92, *D*_*XY*_ 0.0011), and rice ortholog functional genes located in highly divergent regions are marked on the respective chromosomes.

We identified 1,021 transposable elements (TE) that have polymorphism between AS and ES population, including 732 deletions and 289 insertions across all individuals (Fig. S7, DataSet S3). Besides, 13,811 structure variants (SVs) were also identified (Table S12). All of these genetic changes were found to occur sparsely across the total genome without linked changes. A neighbor-joining tree and a PCA based, respectively, on SV data and transposable elements polymorphism (TEPs) also clearly distinguished the AS and ES populations from each other (Fig. S8), indicating that their divergence across the total genome may have involved ecological adaption.

To further investigate the origins of the *B. stacei* AS and ES population at ECI we phylogenetically analyzed the maternally inherited plastomes of this species including samples from ECI, other regions of Israel, and other native Mediterranean locations. A maximum likelihood plastome gene tree rooted with *B. distachyon* Bd30 supported the strong monophyly of the ECI group (tandem plastome gene**s** tree, Fig. S9A; full plastome tree, Fig. S9B), which was nested within a strongly supported ECI-Israel p. p. (*pro partim*) clade (Fig. S9). Within the ECI lineage, all ES samples were shown to be derived from one of the AS lineages (Fig. S9B) and a parsimony network of plastome haplotypes from the two slopes provided further support for this scenario (Fig 2B).

### Demographic divergence of two *B. stacei* populations in ECI

To reconstruct the most plausible evolutionary scenario of divergence between AS and ES populations of *B. stacei* in ECI, we simulated alternative demographic history models of the two populations using forward simulation and residuals analysis in *∂a∂i*. We tested seven models by fitting a site frequency spectrum (SFS) of the AS and ES populations (Fig. 2C, S10A). The best fitting model (based on likelihood and AIC values) suggested a single population divergence event with different reciprocal and asymmetric migration rates in two different time spans (T1 and T2; “asym_mig_twoepoch” model) (Fig. 2C, S10B, Table S13, S14). According to this model, the most recent common ancestor of the AS and ES populations split ∼10.16 kya and had an estimated population size of 13,421 individuals. The first epoch (T1) lasted to the start of the second epoch (T2) ∼3.04 kya, which lasted until the present, and the AS and ES populations had estimated populations sizes of 4,505 and 2,654 individuals, respectively (Fig. 2C, Table S13). According to the model there were continuous and reciprocally asymmetric migrations between both populations in both time periods, with lower migration rates in the second period. These results suggest that continuous gene flow has occurred between the two populations, although gene flow decreased with divergence time (Fig. 2C, Table S14).

A similar distribution pattern of population nucleotide diversity (π) in the AS population (mean π = 1.7893 × 10^−3^) and ES population (mean π = 1.7810 × 10^−3^) was observed (*p* = 0.861, t-test) (Fig. S11), indicating that both populations maintained similar genetic diversity in their respective habitats. In addition, we detected similar genome-wide linkage disequilibrium (LD, indicated by *r2* values) between individual genomes of the AS and ES populations (Fig. S12A). Half of the maximum *r2* value indicated an average physical distance between SNPs of ∼200 kb in the genomes of each population (Fig. S12A), a much longer LD decay distance compared with reported distances of other species (23-25). We also corroborated the high inbreeding rate of the *B. stacei* AS and ES ECI populations (inbreeding coefficient>0.84; selfing rate>0.92; Fig. S12B), as reported for other circum-Mediterranean populations based on molecular markers (26).

The estimated average genome-wide genetic divergence between the AS and ES populations, expressed in terms of fixation index (*F*_ST_) was ∼0.336 (Fig. 2D, Fig. S13). We detected some highly divergent regions (*F*_ST_ ≈ 1) between AS and ES populations (Fig. S13C, Dataset S4). We combined *F*_ST_ and genetic divergence (*D*_XY_) values to identify regions of the genome that were resistant to gene flow (Fig. 2D, Fig. S14). A total of 915 genes were identified from 6.06 Mb highly divergence genome regions at all chromosomes across the total genome (Fig. 2D, Dataset S5). We then explored whether these divergent alleles formed ‘genomic island’ based on the top 5% of the *F*_ST_ and *D*_XY_ values through linked sweeping. We only recovered 3.05Mb sequence of genome, which contained 5 ‘islands’ >70kb, 20 of 40-70kb and 100 of <=30kb (Table S15), and three class islands scattered in all chromosomes (P-value = 0.4733, Fisher’s exact test).

Some rice orthologs of these genes were found to be functionally related to reproduction (*UMA3, EPAD1, TUB8*), plant-pathogen interactions (*NDPK1, NPR3, RacGEF1*), responses to abiotic stress (*Gnk2RLK-1, ZFP182, SIT1, SRWD2, IRT1*), and cell-cycling (*CDKA2*) (Fig 2D). As a complement to the *F*_ST_ and *D*_XY_ approaches, we also applied Hudson-Kreitman-Aguadé (HKA) tests to identify genes under recent selection. We found that 546 genes (Dataset S6) involved in plants’ responses to abiotic stress were significantly enriched (Fig. S15, Dataset S7). For example, we discovered three fixed SNPs in the coding regions of *PIN3A* between AS and ES populations, two of which were non-synonymous coding mutation sites. A homolog of this gene encodes a putative auxin efflux carrier in rice, and its over-expression can improve the drought tolerance of rice (27). In addition, rice orthologs of the genes *Sta2* (28), *HsfB2b* (29), and *PEX11-4* (30) reportedly contribute to salt or drought responses. The AS and ES polymorphisms in these abiotic stress response genes may reflect selective adaptation to their respective microhabitats. These analyses suggest that the genomic divergence between the two *B. stacei* populations from opposite slopes of ECI resulted from ongoing local adaptation to contrasting microhabitats, which may lead to incipient sympatric speciation in these populations.

### Transcriptomic and metabolomic divergence between AS and ES populations of *B. stacei*

To further elucidate potential mechanisms of local adaptation in AS and ES populations in ECI, we applied multiple-level comparisons. We first conducted drought experiments by growing individuals from AS and ES populations in both well-watered and drought conditions to simulate the contrasting ECI biomes. The above-ground plant phenotypes of AS and ES individuals were similar under well-watered conditions, but differed slightly under the drought treatment (Fig. 3A). Five physiological parameters differed significantly between plants grown in drought and well-watered conditions; transpiration rate (E), assimilation rate (A), intracellular carbon dioxide concentration (Ci), and stomatal conductance (Gsw) values were lower, while water use efficiency (WUE) values were higher, under drought than under watered conditions (Fig. S16). AS and ES individuals showed similar changing trends although the AS plants showed higher values in most cases.

**Figure 3.**
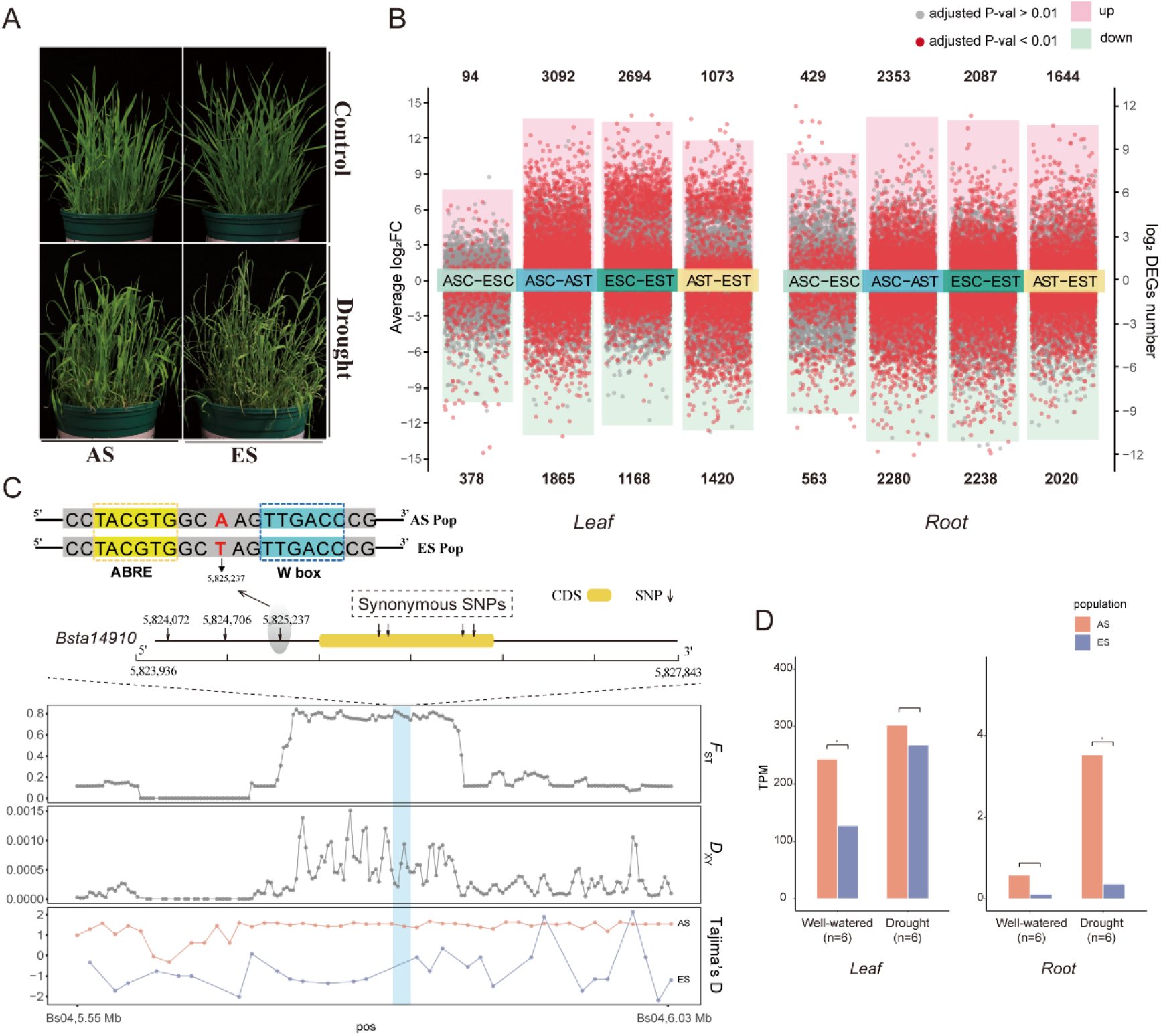
Phenotypic, transcriptomic, and genetic responses related to drought stress conditions of *Brachypodium stacei* plants from AS and ES populations. **A)** Above-ground phenotypes of plants from AS and ES populations grown under drought and well-watered (control) conditions. **B)** Up-regulated and down-regulated differentially expressed genes (DEGs) in leaf and root tissues from intra- and inter-AS and ES population comparisons; adjusted p-values < 0.01 and > 0.01 are indicated by red and gray dots, respectively. **C)** Genetic structure and SNP variants located between +/-2 kbp (upstream and downstream) of gene *Bsta14910* (ortholog of rice OsNCED1) in chromosome Bs04. Population parameters *F*_*ST*_, *D*_*XY*,_ and Tajima’s D of AS and ES populations. The bright blue bar in the plots represents the windows containing *Bsta14910* and its polymorphisms. **D)** Comparative differential expression of *Bsta14910* in leaf and root tissues of AS and ES individuals under control (well-watered) vs treatment (drought) conditions. Values represent transcripts per million (TPM), *p-value < 0.05 (Wilcoxon rank sum test).

To identify key genes in drought responses, RNA-seq analysis was applied and Pearson correlation coefficients showed good repeatability of gene expression profiles among replicates from each population (Fig. S17). We compared gene expression levels in leaf and root tissues from AS and ES plants grown under control (well-watered) and drought conditions, in both inter- and intra-population comparisons (Fig. 3B). In agreement with the observed phenotypic and physiological differences, numerous differentially expressed genes (DEGs) were detected between plants grown in the contrasting conditions in the intra-population comparisons (Fig. 3B). In total, 3,862 and 4,325 DEGs were identified in leaves and roots of ES individuals, and more (4,957 and 4,633 DEGs) in those of AS plants (Fig. 3B). Diverse Gene Ontology (GO) terms and KEGG pathways related to drought responses were enriched in sets of these DEGS in both AS and ES plants (Fig. S18, Dataset S8). 1,630 common DEGs in leaves and 1,423 common DEGs in roots were identified in inter-population comparisons between AS and ES plants (Fig. S19). The Log_2_ fold change (Lfc) values of these differentially expressed genes were similar for the AS and ES plants (Pearson correlation, R≥ 0.75, *p*< 2.2e^-16^) (Fig. S20), suggesting that drought stress induced similar qualitative effects on the gene expression of leaf and root in the AS and ES individuals although their respective expression levels were significantly different.

Only 462 leaf DEGs and 992 root DEGs were identified from AS-ES inter-population comparisons under well-watered conditions (Fig. 3B), suggesting that gene expression is quite similar in both populations under well-watered conditions. In contrast, 2,493 leaf and 3,664 root DEGs were identified from AS-ES inter-population comparisons under drought conditions, 5.3-fold and 3.6-fold higher than numbers identified from corresponding comparisons of well-watered plants (Fig. 3B), suggesting potential differences in drought response mechanisms between individuals of the AS and ES populations. The observed dissimilarities in genome and transcriptome between AS and ES populations were mirrored by their respective metabolomes. We analyzed metabolite profiles of leaf and root tissues from AS and ES plants under drought conditions using Liquid Chromatography - Mass Spectrometry (LC-MS). In inter-population metabolome comparisons, 158 and 120 differential metabolites (DM) were identified in the leaf and root tissues, respectively. Orthogonal partial least-squares discriminant analysis (OPLS-DA) separated the AS and ES metabolomic profiles into two clusters, indicating that the metabolite profiles of AS and ES plants grown under drought conditions clearly differed (Fig. S21, Datasets S8). The DM data were consistent with the finding that KEGG biological pathways related to drought stress responses were significantly enriched in the DEGs of the AS and ES populations (Fig. S22, Datasets S8, S9). We further investigated the expression of genes directly related to drought and oxidative responses, several of which had different expression patterns in the AS and ES populations, for example, *DRO1, APX8*, and *ECK1* (Fig. S23). These different combinations of gene expression and metabolite data suggest that AS and ES *B. stacei* populations respond discordantly to drought stress, likely as a consequence of distinct local adaptations to their contrasting microclimates.

### Key factors regulating microclimate adaptation in *B. stacei* ECI populations

We investigated genes involved in the abscisic acid (ABA) signaling pathway with differing expression patterns in the AS and ES populations, as it is one of the main regulatory systems of plants, with broad effects, especially in abiotic stress responses. Catalysis of the transformation of 9-*cis*epoxycarotenoids to xanthoxin, by 9-*cis*-epoxycarotenoid dioxygenases (NCEDs), is a key regulatory step in ABA biosynthesis (31). We found that a putative ortholog of rice *OsNCED1* (*B. staceiBsta14910*) (Table S16) showed high genetic divergence and expression changes between the AS and ES populations (Figs. 3C, 3D). In total, four SNPs located in the coding region of this gene corresponded to reciprocal synonymous amino acid mutations between the two populations (Fig. 3C). We also found 16 SNPs in the 5’-upstream 2000 bp regulatory region of *Bsta13013*, all of which detected a high divergence between the two populations (Table S17). Three SNPs were fixed in the AS and ES populations, including one located 699 bp upstream of the start codon (Fig. 3C). Further analysis showed that this SNP was located between the TACGTG (ABRE) and TTGACC (W box) motifs (Fig. 3C), suggesting that these mutations may affect the transcription of *Bsta14910*, which was significantly overexpressed in leaf and root tissues of the AS samples compared to the ES samples (Fig. 3D). Other *NCED1* orthologs are involved in heat responses in rice (32) and *Lactuca sativa* (33), underscoring this gene’s potential importance for adaptation to warm conditions.

We also found that exposure to drought induced stronger changes in the AS population than the ES population samples in expression of a gene encoding a serine/threonine protein kinase (*SAPK5*), which plays a key role in ABA signaling pathways activated by hyperosmotic stress (34) (Fig. S24). In addition, we identified a *Glossy1 B. stacei* ortholog (*GL1-2*) with significantly higher expression patterns in the AS than ES populations samples (Fig. S25). This gene is induced by ABA and reportedly involved in wax biosynthesis, and thus may participate in important adjustments of the composition of leaf waxes that enhance resistance to abiotic stressors, such as drought, ultraviolet light, and extreme temperatures (35). Together, these results suggest that differential expression of genes encoding key proteins involved in ABA signaling and wax synthesis may play important roles in different adaptations to local environments in the AS and ES populations. In addition, the changes of TEs may affect the expression of nearby gene through altering or creating regulatory element during ecological divergence (23). We identified 15 TE insertion polymorphisms (TIPs) that have different frequency between the two populations (Fig S7; DataSet S9) and nine genes were associated with them. Two genes (*Bsta12548* and *Bsta19483*) showed stable differential expressions between the two populations, being more highly expressed in AS than in ES samples in both tissues and conditions (Fig. S26). *Bsta12548* is involved in post-translational modification and protein turnover process while *Bsta19483* encodes a wall-associated receptor kinase (35). Therefore, mutations in regulatory motifs and coding regions of such genes and their targeting genes may also contribute to disconnected local adaptations and genetic divergence in the arid AS and mesic ES populations at ECI.

## Discussion

Ecological divergence has been frequently reported for organisms living in ECI (see Dataset 10) because of the canyon’s striking differences in microclimate and biomes (7, 8, 36). Diverse organisms, from bacteria to mammals and plants, have undergone ecological divergence in ECI, where the contrasting ecological conditions have promoted prezygotic and/or postzygotic reproductive isolation that may have led to the emergence of different species (8, 18, 37-39). Populations of some species (e.g., wild emmer wheat and spiny mouse) on the two slopes of ECI have been reproductively isolated through prezygotic (14, 18) (including differences in flowering time in plants and mate discrimination in animals) and postzygotic (19) isolation mechanisms (including chromosome re-arrangements (18) although not completely. In this study, we found that the ECI’s contrasting microclimates have similarly fostered genomic, transcriptomic, and metabolomic divergence between two diploid inbreeding *B. stacei* populations at ECI (Figs. 2 and 3), suggesting ongoing divergence and further possible production of two species with complete reproductive isolation in the future.

Our population analysis of phylogenetically inherited plastomes showed that both AS and ES individuals shared a common maternal ancestor compared to other *B. stacei* accessions out of ECI (Fig. 2B, Fig S9), supporting that the two populations diverged *in situ* although we could not totally rule out the possibility that they diverged elsewhere and migrated to and survived at ECI. In addition, we found that ES individuals formed a sub-clade nested within the AS clade in a strongly supported ML plastome tree, suggesting that *B. stacei* established first in the AS and then later colonized the ES (Fig. S9). The lower divergence of recently expanded ES individuals compared to the larger divergence of more ancestral AS individuals was also supported by all nuclear genomic components (SNPs, SVs, TEPs; Figs. S6A, S8A, S8B). In previous evolutionary studies, both colonization scenarios from AS to ES and from ES to AS have been found (9, 11, 14, 18, 40-42), but our plastome data clearly support an AS to ES colonization scenario for *B. stacei* in ECI (Fig. 2B, Fig S9). Despite the non-reciprocally monophyletic clustering of individuals from each slope in the plastome tree, which is likely a reflection of colonization history, nuclear genome data also differentiated the two populations (Figs. 2A, S6, S8), suggesting that individuals of each population share their own genome variants. All of these analyses demonstrate that both populations have evolved as separate independent lineages.

Our demographic analyses suggest that the AS and ES populations of *B*. stacei diverged approximately 10,000 years ago, and gene flow gradually decreased with increasing divergence time (Fig. 2C, Fig. S10, Tables S13, S14). This finding supports the prediction that gradual divergence of two populations in contrasted habitats may lead to sympatric speciation (43). We found evidence of genetic divergence (mean *F*_ST_ = 0.33) between the two populations that is higher than the genetic divergence observed in other plant and animal species that have undergone sympatric divergence at this site (11, 18). This may have resulted from the high inbreeding rates of *B. stacei* that promote selective fixture during ecological stress (Fig. S12B) (26, 44). Our results further suggest that gene flow between AS and ES populations is mainly mediated by seed dispersal although inter-population pollen exchange may still occur in rare cases (Fig. 2B; Fig. S6B). Despite the continuous gene flow, we provided the clear evidence that the two populations of *B. stacei* sympatrically diverged in response to disruptive selection pressures on the two slopes (43, 45). Individuals from the AS population have significantly more drought tolerance than those from the ES population, as evidenced by transcriptomic, metabolomic, and physiological trait analyses (Fig. 3, Figs. S16-S26). Functional genomic (46) and metabolomics (47) studies of the closely related species, *B. distachyon*, also found significantly greater induction of genes and metabolites important for drought stress responses in individuals adapted to arid conditions compared to those adapted to mesic conditions. These differences at multiple loci probably maintain sympatric separation of the two populations by facilitating divergent local adaptation to contrasting habitats.

The highly diverged and fixed in both coding and noncoding (TEs) alleles in frequencies between two populations were revealed to be distributed across all chromosomes of the *B. stacei* genome (Table S15, DataSet3). Only very few genes formed small ‘genomic islands’ through sweeping links, which is consistent with the experimental test of genomic divergence under ecological divergence with continuous and strong gene flow (4). However, this contrasts with the ecological divergence of a few species in the sympatric site in which the large-scale linked genomic divergences comprised the obvious ‘genomic islands’ within a few chromosomes (5) possibly due to micro-parapatric speciation with partly geographic isolation, or the second contacts after the initial allopatric divergence in other regions. In conclusion, our study provides robust sources from genomic, transcriptomic, and metabolomics analyses for the ecological divergence of two *B. stacei* populations with a most recent ancestor growing on the ECI opposite slopes. In addition, multiple unlinked loci may act additively to contribute to such an ecological divergence. These cumulative evidences support the initial sympatric speciation scenario of *B. stacei* in ECI as indicated by other studies (4, 48, 49).

## Materials and methods

The genome was sequenced using a PacBio Sequel2 platform and assembled using NextDenovo. The whole-genome DNA re-sequencing data were generated by an Illumina HiSeq X Ten machine. Multiple alignment files were generated with BWA-MEM2 (v2.2.1). Population structure was analyzed using ADMIXTURE (v1.3.0). Phylogeny trees was constructed with IQ-TREE (v 1.6.12). *F*_ST_ and *D*_XY_ were calculated by Pixy. Transcriptome analysis was conducted using the ‘HISAT2-Stringtie-DESeq’ pipeline. Detailed experiments and analyses are available in SI Appendix.

## Data availability

The sequencing data are deposited at NCBI, the project number are PRJNA791186 and PRJNA791713, which can be accessed through URLs (https://dataview.ncbi.nlm.nih.gov/object/PRJNA791186?reviewer=ivdf940cv986vs92hvu907vh07). The genome assembly and main script used in the analyses have been uploaded at Github (https://github.com/Axolotl233/Brachypodium_SS)

## Author contributions

P.C. J.L., K.L., and E.N. designed the study. P.C. K.L. and A.B. conducted the sampling.

W.M. and Y.Y. performed the experiments and the analyses. J.Y., Y.W., and M.Z. contributed to the analyses. P.C., J.L., E.N., W.M and wrote the manuscript. All authors read and commented on the manuscript.

## Acknowledgements

All the computation works were supported by the Big Data Computing Platform for Western Ecological Environment and Regional Development, and the Supercomputing Center of Lanzhou University.

