## Supplemental Figures 1-26 and Table 1-17 for "Scattered differentiation of unlinked loci across the genome underlines ecological divergence of the selfing grass *Brachypodium stacei*"

### **Supplementary Figures: *Brachypodium stacei***

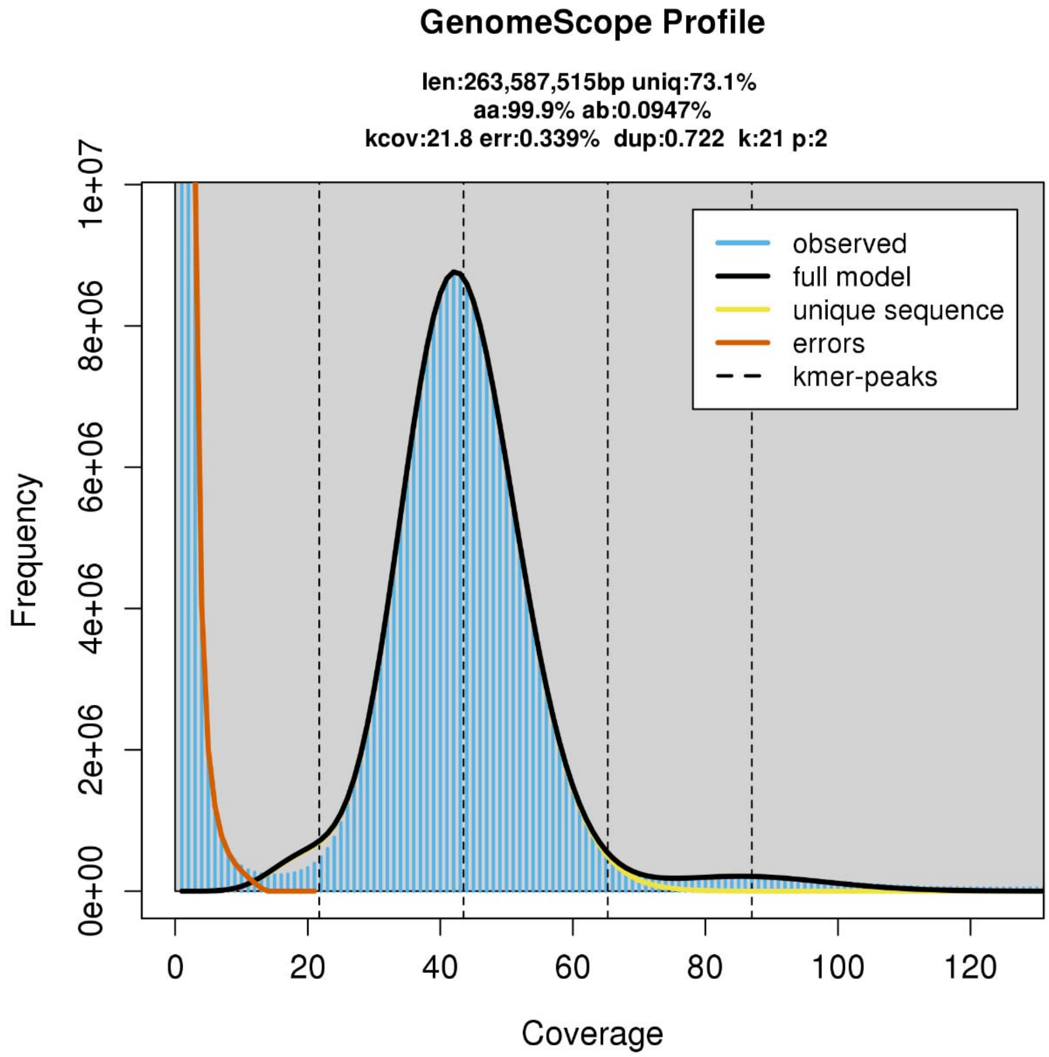


**Figure S1**. Genome survey of the *Brachypodium stacei* Bsta-ECI accession from Evolution Canyon I (see Supplementary Table S1 for additional information)**.** K-mer size was set at 21 and the default parameters were set with Genomoscope2.

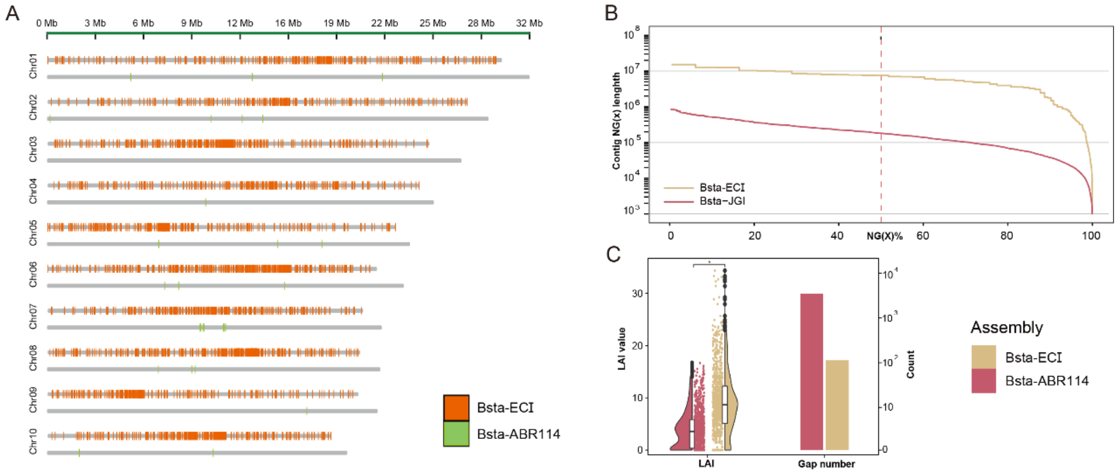


**Figure S2.** Compared genomic features of the new *B. stacei* ECI assembly from Evolution Canyon I and *B. stacei* reference genome ABR114 obtained from Illumina sequencing at the Joint Genome Institute (Phytozome: https://phytozome-next.jgi.doe.gov/). **A)** Chromosomes length statistics of genome assembly, gap positions with respect to genomes were marked as color bars.. **B)** Overview of assembly contig lengths comparing NG values , from 1 to 100%, and the contig length (in bp) for particular thresholds are shown on the y-axis. Calculated NG contig lengths range on a log scale. The dashed vertical line indicates the NG50 contig length. **C)** Comparison of gap numbers and the LTR Assembly Index (LAI) among assembly versions, *p-value < 0.01 (t-test).

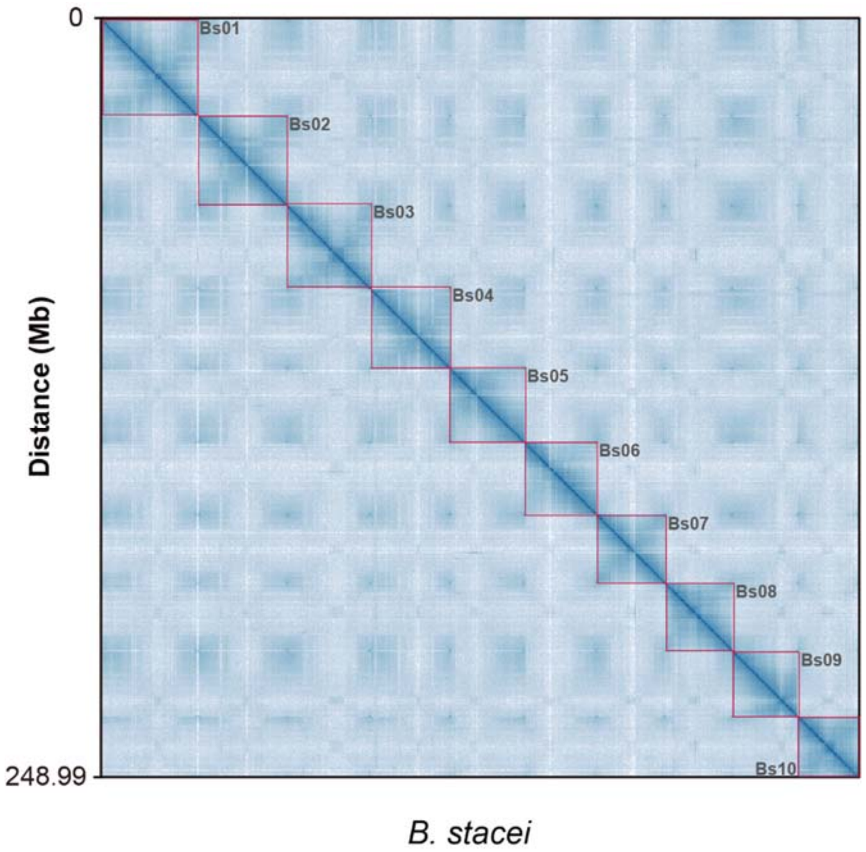


**Figure S3.** Newly assembled *Brachypodium stacei* ECI genome. Hi-C interaction densities heatmap between contigs; its 10 chromosomes (Bs01-Bs10) are separated by red boxes.

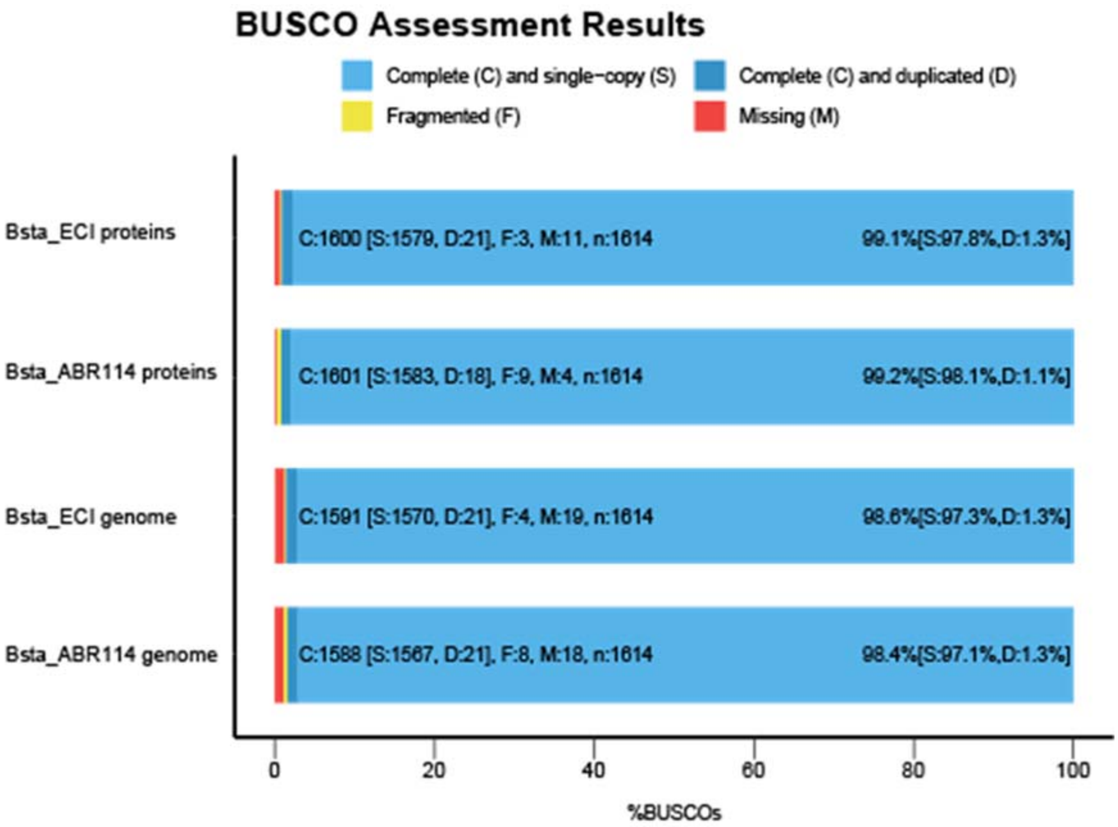


**Figure S4.** BUSCO gene assessments of genome assemblies. Color codes for gene types are indicated in the chart. Bsta-ECI, newly assembled *Brachypodium stacei* genome from Evolution Canyon I; Bsta-ABR114, *B. stacei* reference genome (Phytozome; https://phytozome-next.jgi.doe.gov/). BUSCO runs in "genome" and "proteins" modes.

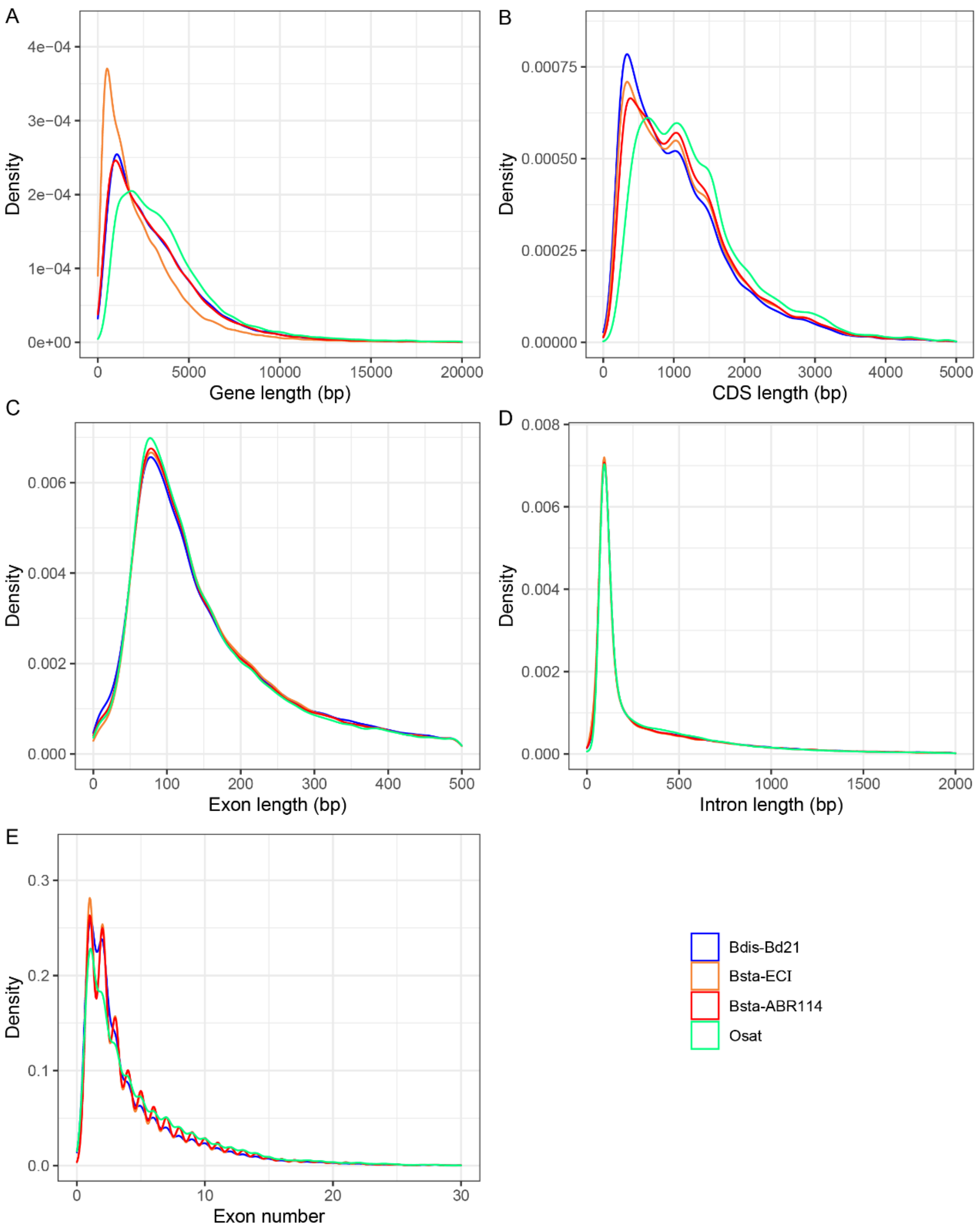


**Figure S5.** Comparison of gene model structure characteristics in the newly assembled *B. stacei* (Bsta-ECI) genome to those in other assemblies (reference genomes of *B. stacei* ABR114 v. 1.1, *B. distachyon* Bd21, generated by the Joint Genome Institute (JGI), see Gordon et al. 2020) and *Oryza sativa* (Osat). (A) gene length; (B) CDS length; (C) exon length; (D) intron length; (E) exon number.

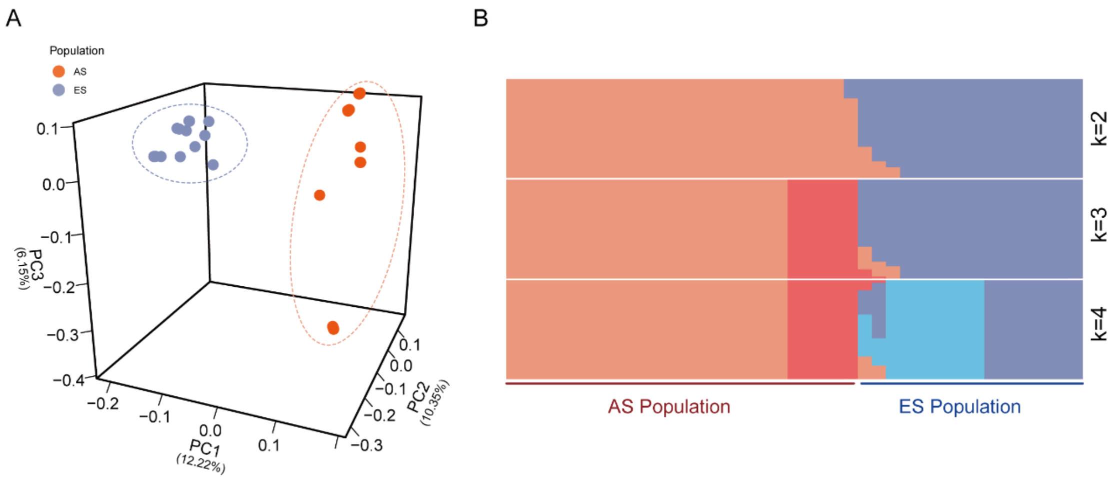


**Figure S6.** Population structure analysis of 41 *Brachypodium stacei* resequenced individuals from Evolution Canyon I (ECI) based on genome SNP data**.** A) Tridimensional plot of Principal Component Analysis (PCA). The PCA1 axis (accumulating 12.22% of the variance) clearly separated the two populations from the African Slope (AS) and the European Slope (ES). B) Population structure bar plot retrieved from Structure for *K*= 2, 3, and 4 hypothetical populations. The optimal structuring corresponded to K=2, which also separated the AS and ES populations; other less-optimal Ks showed some substructuring within AS (K=3) and AS and ES (K=4).

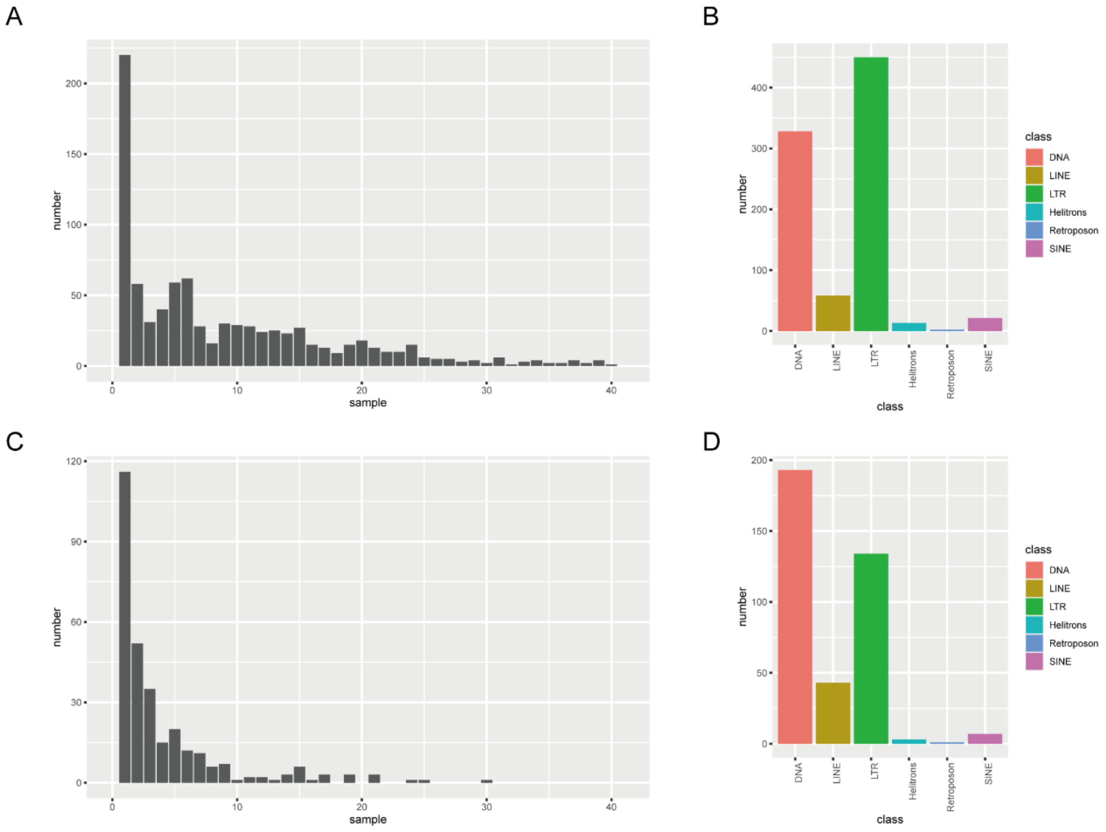


**Figure S7.** Distribution and classification of transposable element polymorphisms (TEPs) found in the 41 *B. stacei* genomes studied from ECI. (A) Distribution of deletions among all 41 genomes relative to the Bsta-ECI local reference genome. The x-axis indicates the number of samples showing deletions and the y-axis the number of polymorphic transposable elements. (B) Classification of polymorphic repeat elements showing deletions with respect to the Bsta-ECI genome. (C) Distribution of insertions among all 41 genomes relative to the Bsta-ECI local reference genome; x and y axes indicate the number of samples showing insertions and the number of polymorphic transposable elements, respectively. (D) Classification of polymorphic repeat elements showing insertions with respect to the Bsta-ECI genome.

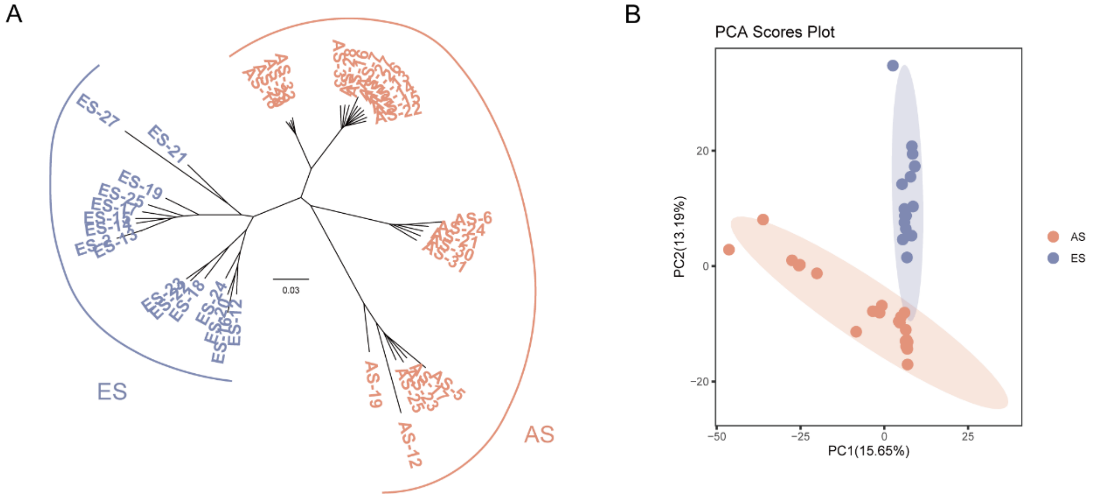


**Figure S8.** Population structure analysis of 41 *Brachypodium stacei* resequenced individuals in ECI using Structure Variants (SVs) and Transposable Elements Polymorphisms (TEPs) data obtained from the 41 resequenced genomes. (A) Neighbor joining tree based on SVs data (see Supplementary Table S12) with variants coded as binary characters; (B) Bidimensional PCA plot based on TEPs coded by frequency classes as binary data. Both analyses separate the samples from the African Slope (AS) and the European Slope (ES) populations.

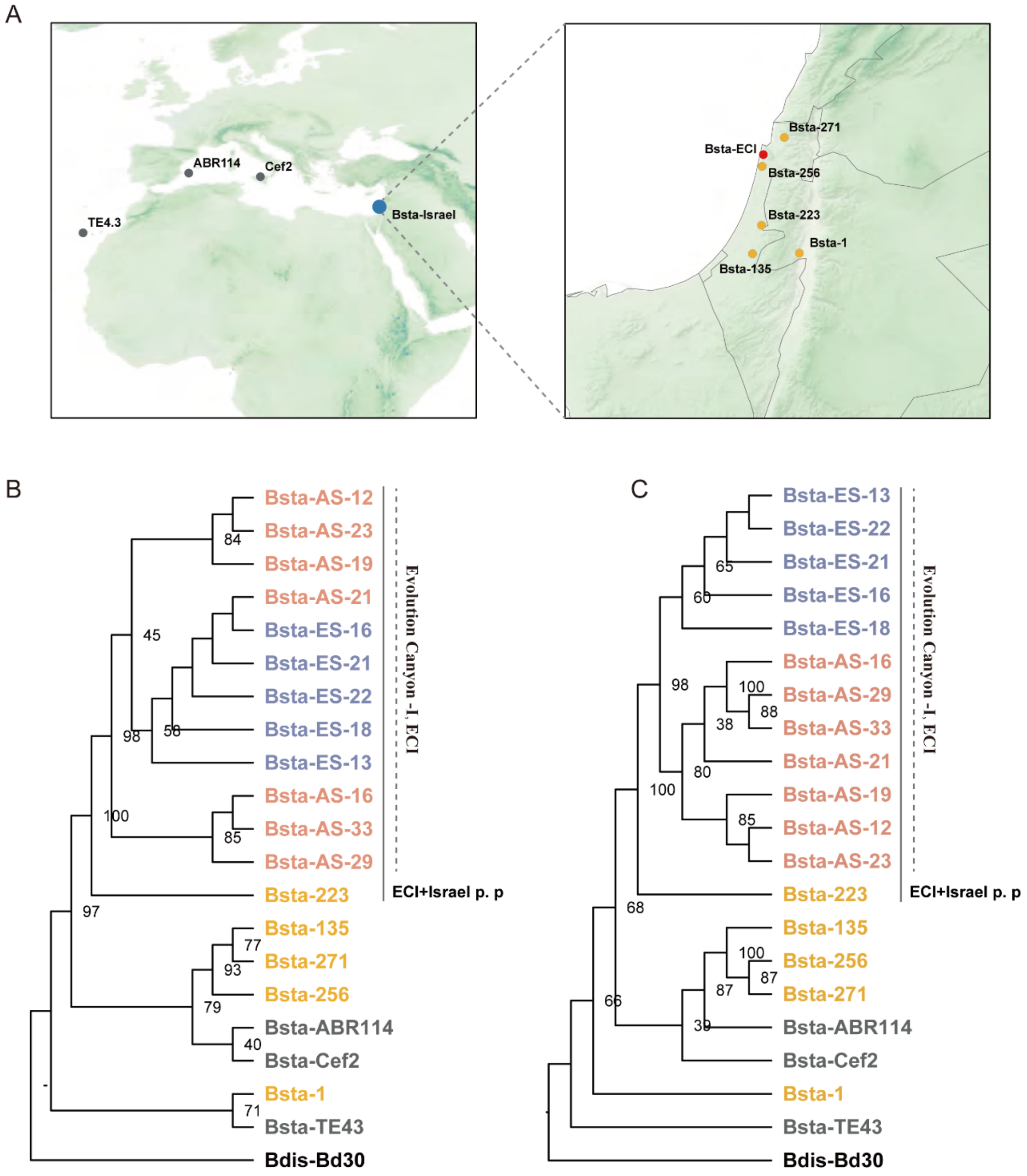


**Figure S9.** The Geographic distribution of *B. stacei* samples used in this study and maximum-likelihood phylogenetic tree based on plastome sequences; numbers at nodes indicate bootstrap support values. **A)** the location of samples **B)** tandem plastome genes tree**. C).** full plastome tree**.** Clades: AS, African Slope; ES, European Slope; ECI, Evolution Canyon I; ECI+Israel p. p. (pro partim), ECI + Is-223.

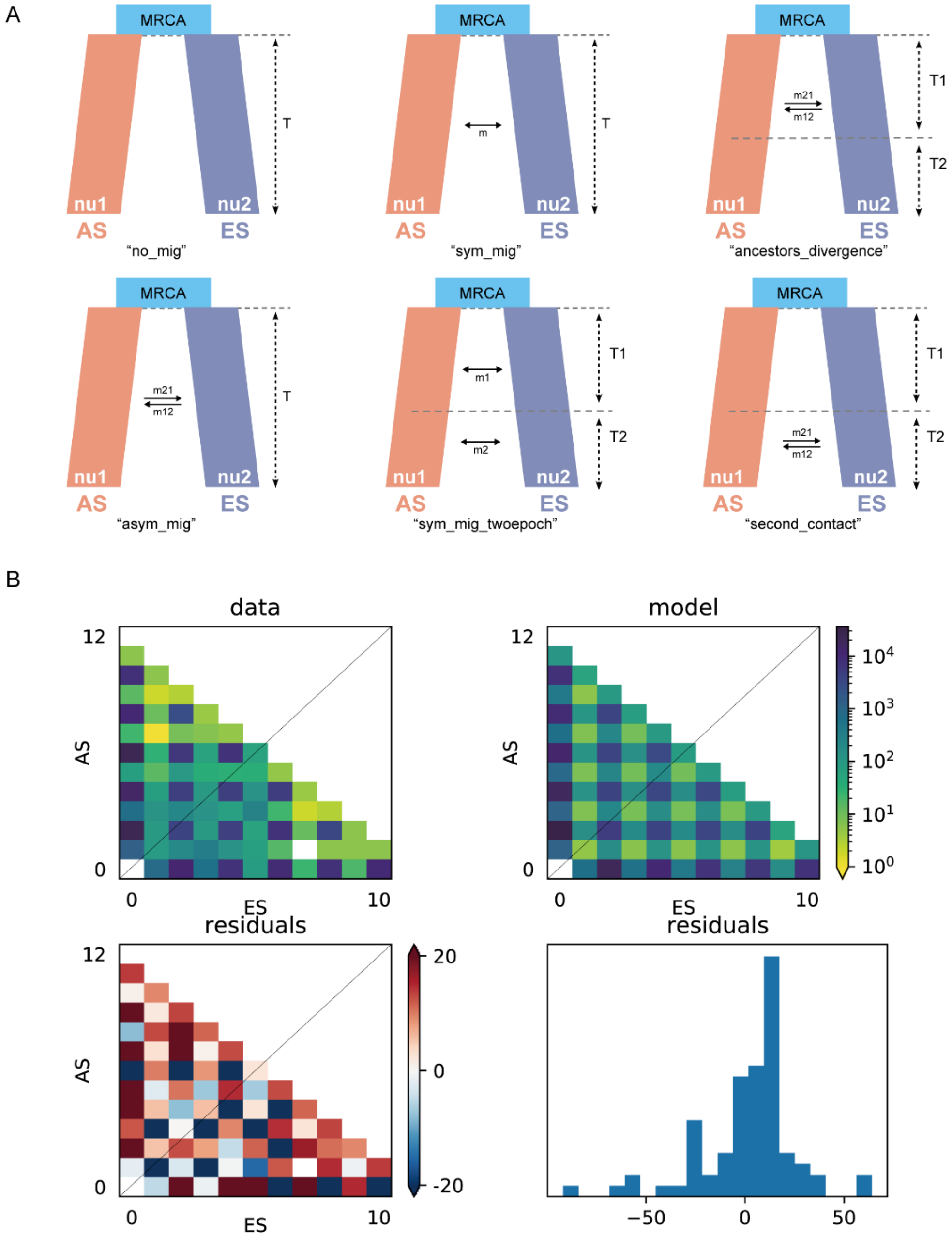


**Figure S10.** (A) Six other alternative *∂a∂i* models used in the inference of the evolutionary demographic histories of the *Brachypodium stacei* AS and ES populations in ECI using forward simulation and residuals analysis. (B) Demographic parameters (p0 = [0.3357; 0.1978; 3.1526; 1.7528; 1.542; 1.3721; 0.1131; 0.3788]) used to perform the residuals analysis of the best fit *∂a∂i* model (see Fig. 2C).

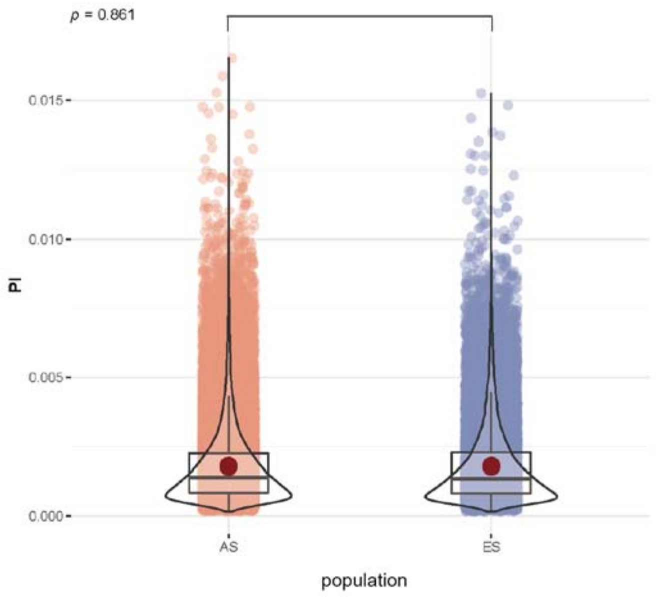


**Figure S11.** Population nucleotide diversity ($\theta_\pi$) of the 41 studied *B. stacei* resequenced genomes at the ECI site. Distribution of $\theta\pi$ values for the AS and ES populations, calculated by 50kb window and 12.5kb step using vcftools. Both populations show similar mean $\theta_\pi$ values.

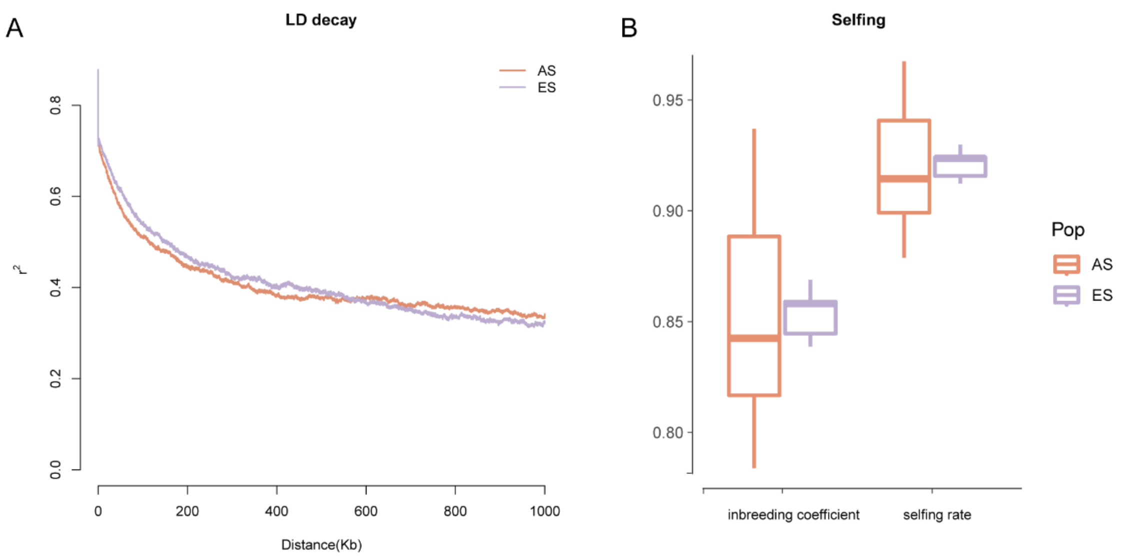


**Figure S12.** Linkage disequilibrium decay (LD) and inbreeding rates of 41 *Brachypodium stacei* resequenced genomes form the African Slope (AS) and European Slope (ES) populations at ECI. (A) LD decay of the two AS and ES populations (x axis: physical distances (Kbp) of the individual genomes; y axis: $r^2$ value); (B) box plots showing inbreeding coefficient values and selfing rate values for the AS and ES populations of ECI calculated with PLINK. Color codes for AS and ES populations are indicated in the charts.

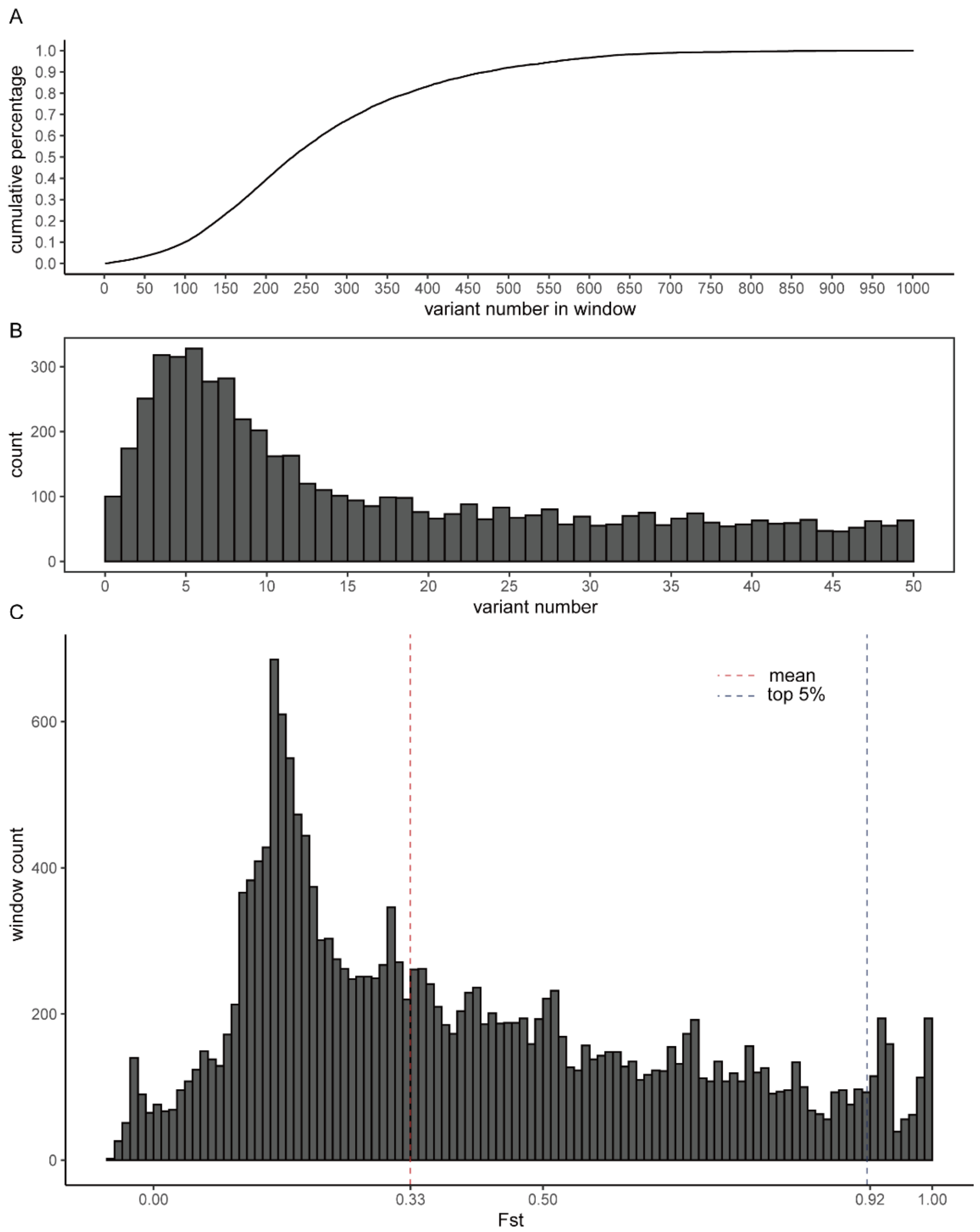


**Figure S13.** Summary statistic of *Fst* values in 50-Kb sliding windows for the *Brachypodium stacei* African Slope (AS) and European Slope (ES) populations at ECI. (A) Cumulative curve of SNP count per window size. (B) Distribution of SNP count in 50-Kb windows, windows containing more than 50 SNPs were omitted from the plot. (C) Distribution of pairwise *Fst* values per number of 50-Kb windows along the whole genome of the studied samples.

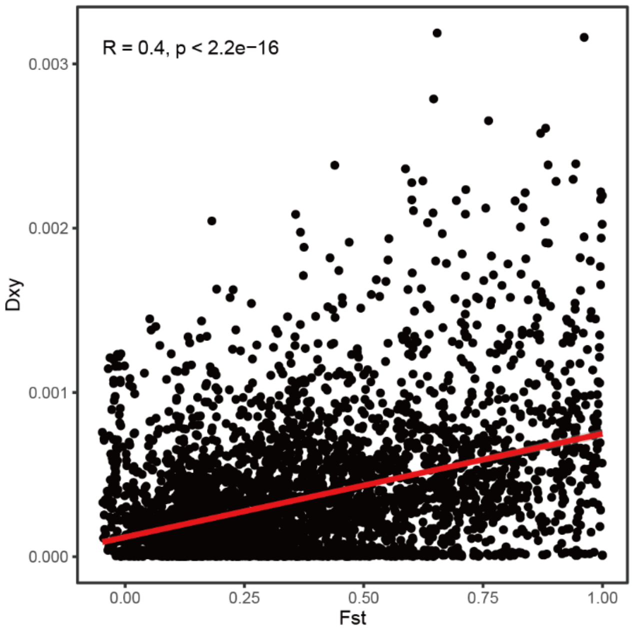


**Figure S14.** Correlation analysis between pairwise $F_{ST}$ fixation values and genetic divergence $D_{XY}$ values between *Brachypodium stacei* AS and ES populations from ECI analyzed across the whole genome (Pearson's correlation test).

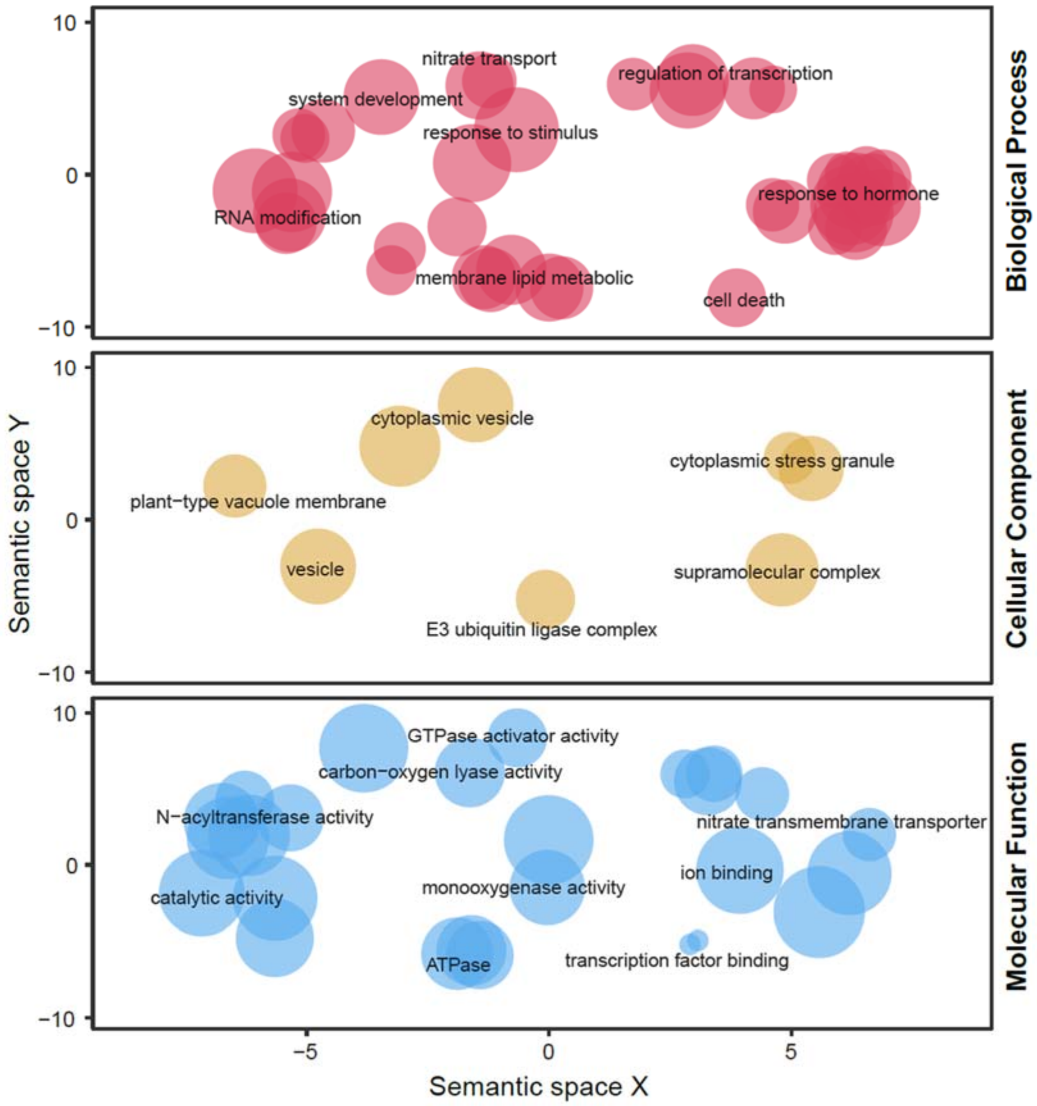


**Figure S15.** REViGO semantic similarity scatter plots of Gene Ontology (GO) terms for *Brachypodium stacei* genes identified by the HKA (Hudson-Kreitman-Aguadé) test as recently selected for Biological processes, Cellular components, and Molecular function. In the semantic spaces, the X and Y axes in the plot represent similarities between GO terms, and the size of the dot indicates gene counts of GO terms.

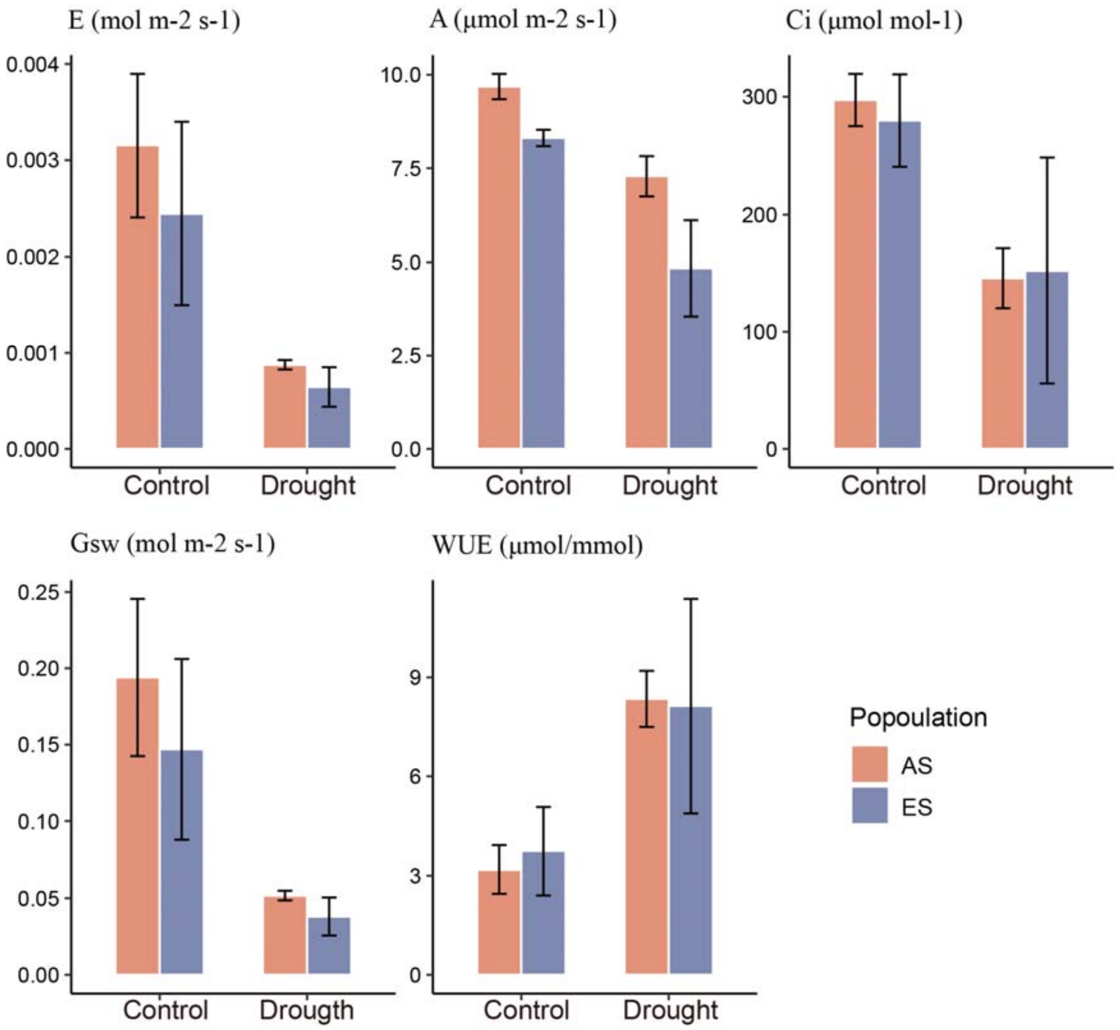


**Figure S16.** Variation of physiological parameters in samples of the two *Brachypodium stacei* AS and ES populations of ECI under treatment (drought) vs control (well-watered) conditions. E, Transpiration rate; A, Assimilation rate; Ci, intracellular carbon dioxide concentration; Gsw, Stomatal conductance to water vapor; WUE, water use efficiency. Color codes for AS and ES populations are indicated in the chart.

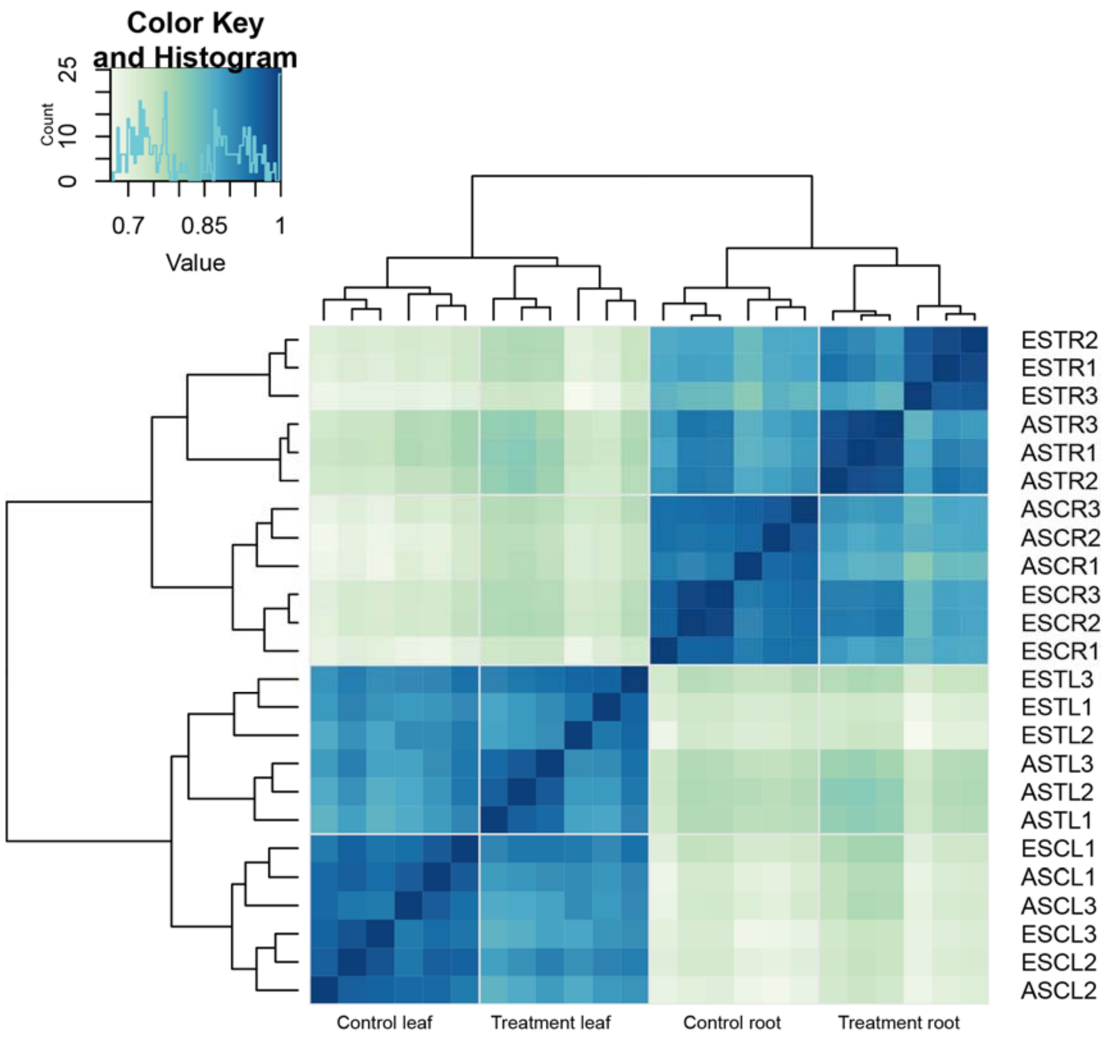


**Figure S17.** A cluster heat map showing the Pearson correlation of the expressed leaf and root genes between *Brachypodium stacei* samples from the African Slope (AS) and European Slope (ES) populations at ECI under drought (treatment) and well-watered (control) conditions. TL: treatment leaf; CL: control leaf; TR: treatment root; CR: control root. Three individual sample replicates from each AS and ES population were used in the expression analysis. Heatmap color code values are indicated in the chart.

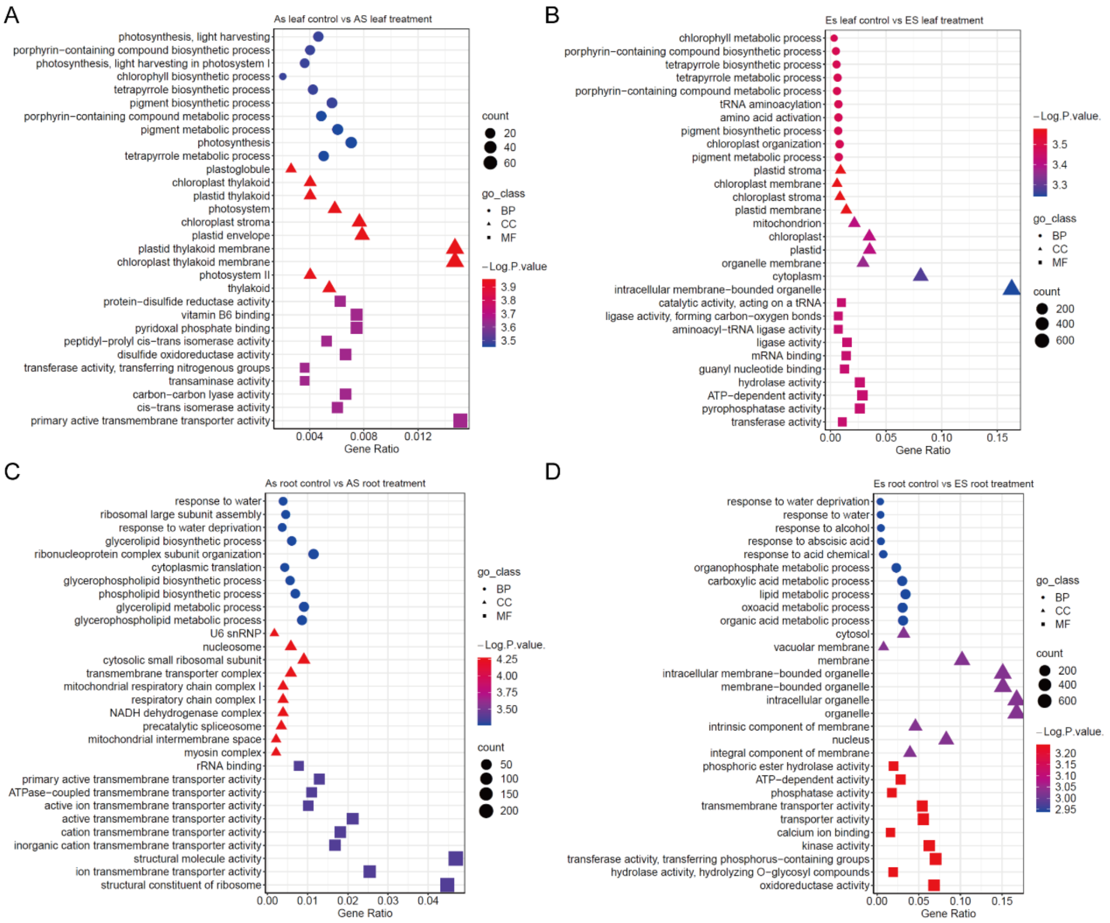


**Figure S18.** Plots of GO enrichment terms of differentially expressed genes in samples of *Brachypodium stacei* from the African Slope (AS) and European Slope (ES) populations at ECI exposed to drought (treatment) *vs* well-watered (control) conditions. Intra-population comparisons, only the top 10 terms are shown. (A) AS leaf control *vs* AS leaf treatment. (B) ES leaf control *vs* ES leaf treatment. (C) AS root control *vs* AS root treatment. (D) ES root control *vs* ES root treatment. Number of genes (circle size and count), GO class and -logPvalue symbols and color codes are indicated in the respective charts.

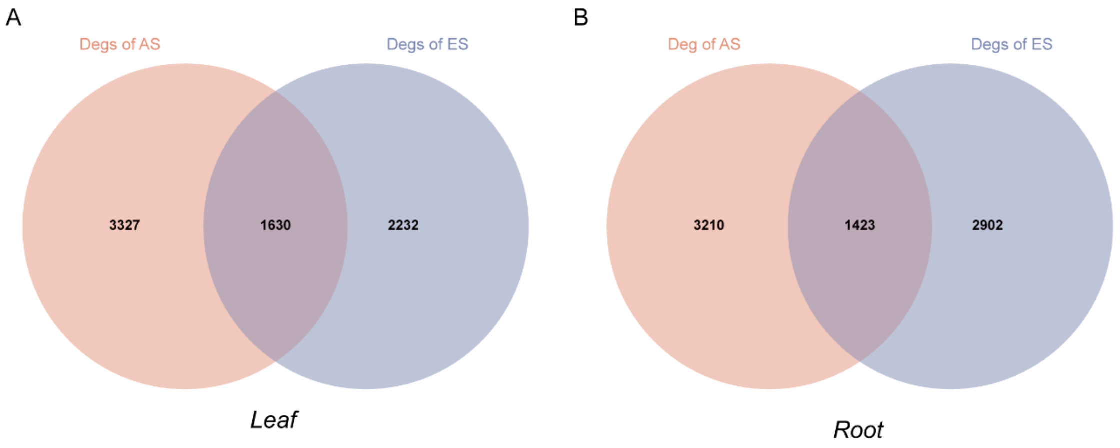


**Figure S19.** Venn plot of differentially expressed genes (DEGs) in inter-population comparisons of *Brachypodium stacei* samples from the African Slope (AS) and European Slope (ES) populations at ECI exposed to drought (treatment) *vs* well-watered (control) conditions. (A) leaf tissue; (B) root tissue.

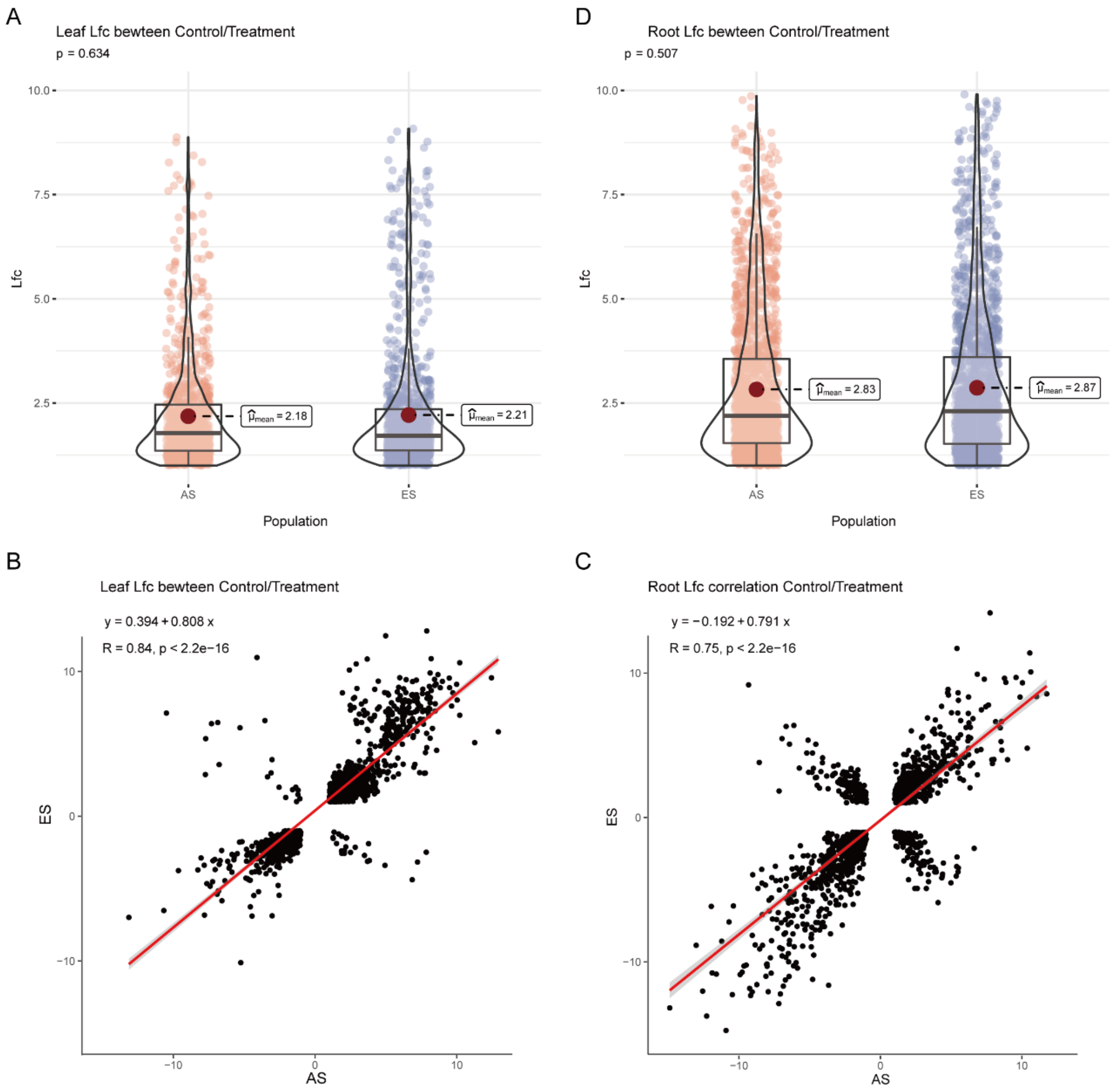


**Figure S20.** Logarithmic fold change (Lfc) distributions of most up and down regulated genes in intra-population comparisons and Pearson correlation coefficients in inter-population comparisons of common DEGs in *Brachypodium stacei* leaf and root tissues of samples from the African Slope (AS) and the European Slope (ES) populations at ECI under well-watered (control) vs drought (treatment) conditions. Only genes with |Lfc|<10 and Pearson correlation coefficient <|15| are shown. (A) Lfcs of leaf DEGs in control *vs* treatment conditions in AS and ES samples (t-test). (B) Lfc of root DEGs in control *vs* treatment conditions in AS and ES samples (t-test). (C) Pearson correlation coefficient of Lcf leaf DEG values from control *vs* treatment conditions (see subfigure A) between AS and ES samples. (D) Pearson correlation coefficient of Lcf root DEG values from control *vs* treatment conditions (see subfigure B) between AS and ES samples. Correlation coefficients were high and significant between the two populations for DEGs' Lfc values in both leaf and root tissues.

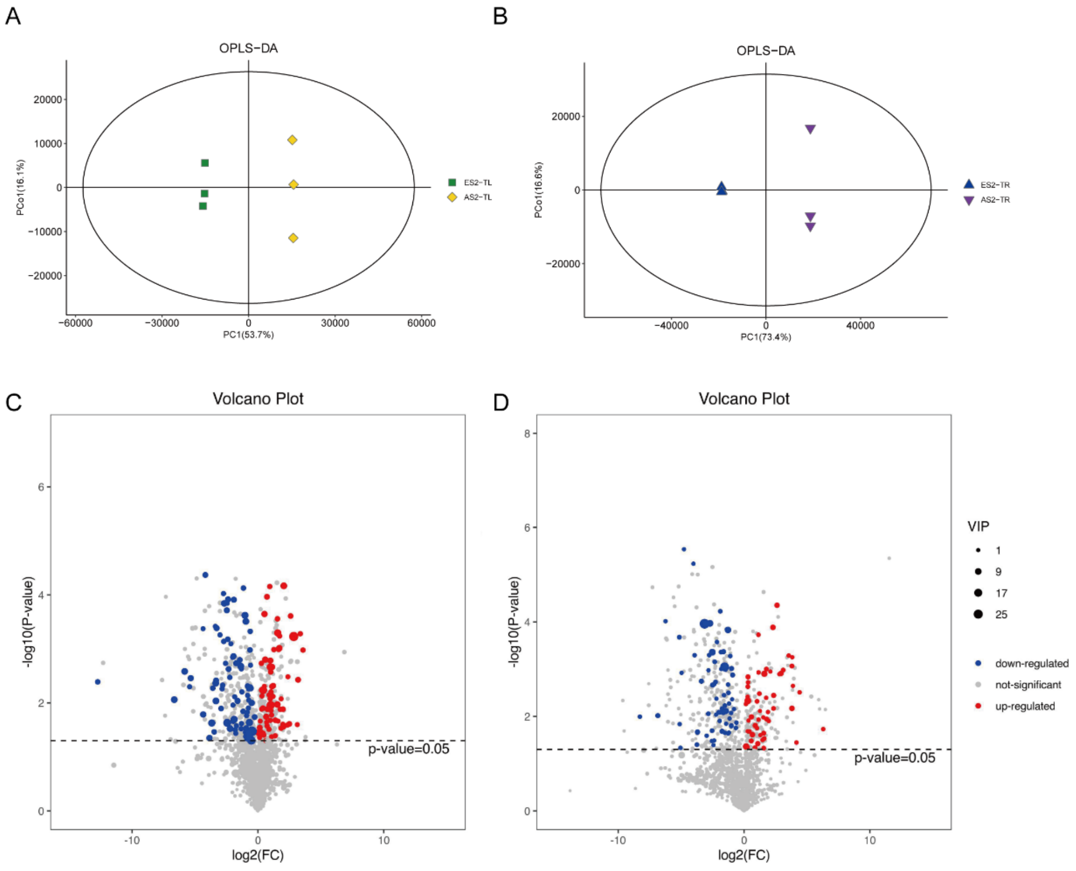


**Figure S21.** Orthogonal partial least-squares discrimination analysis (OPLS-DA) and Variable importance plot (VIP) of metabolomic profiles of *Brachypodium stacei* samples from the African Slope (AS) and European Slope (ES) populations at ECI. (A, B) Bidimensional OPLS-DA plots of metabolite profiles in leaf (A) and root (B) samples under drought conditions from both populations. Symbols and color codes are indicated in the charts. (C, D) Significantly regulated metabolites (up regulated red; down regulated blue) were identified by |FoldChange| > 1 in leaf (C) and root (D) tissues of the studied populations' samples under drought treatment. TL: (drought) treatment leaf; TR: (drought) treatment root. VIP codes (circle size) and regulation codes (colors) are indicated in the chart.

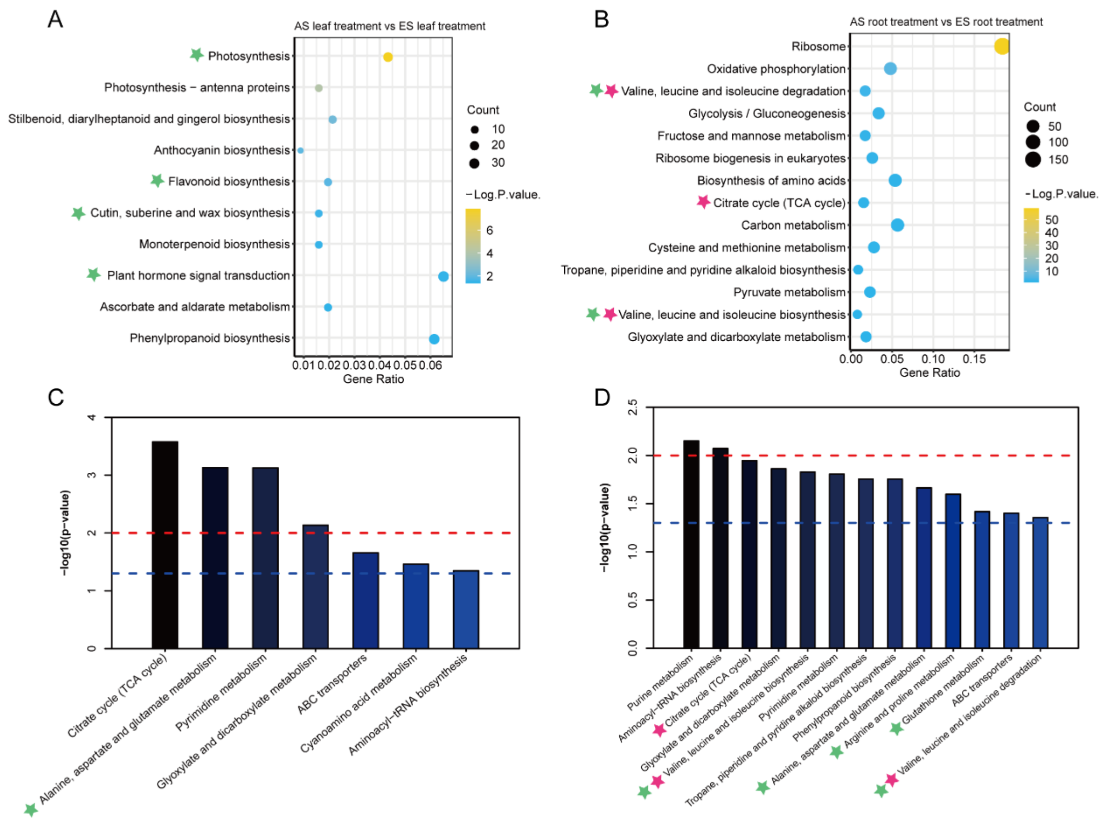


**Figure S22.** Comparative analysis of enriched transcriptome and metabolome KEGG pathways of *Brachypodium stacei* AS and ES population samples from inter-population comparison under drought stress treatment (AS treatment vs ES treatment) for leaf and root tissue samples. Terms involved in drought stress response are marked with a green star and those enriched in both transcriptome and metabolome analysis are marked with a red star. (A, B) Enriched transcriptomes: (A) DEGs enrichment of AS *vs* ES leaf samples under drought conditions. (B) DEGs enrichment of AS vs ES root samples under drought conditions. Gene count (circle sizes) and –LogPvalue (color codes) are indicated in the respective charts. (C, D) Enriched metabolomes: (C) Differentially regulated metabolites enrichment of AS vs ES leaf samples under drought conditions. (D) Differentially regulated metabolites enrichment of AS vs ES root samples under drought conditions. Dashed blue and red lines indicate p-values < 0.05 and <0.01, respectively.

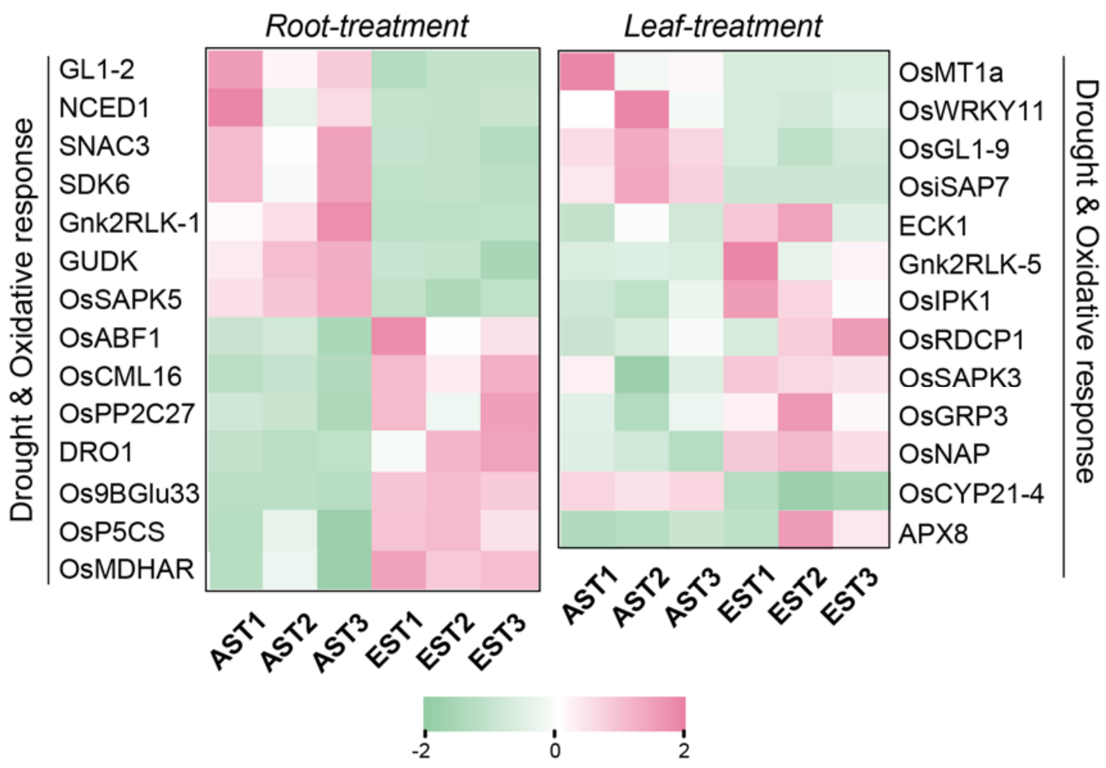


**Figure S23.** Heatmap of *Brachypodium stacei* DEGs for drought response and oxidative response in root and leaf tissues from AS and ES samples under drought (treatment) conditions. AS and ES root and leaf samples consisted of three biological replicates each (T1-T3). Color codes of the heatmap are indicated in the chart.

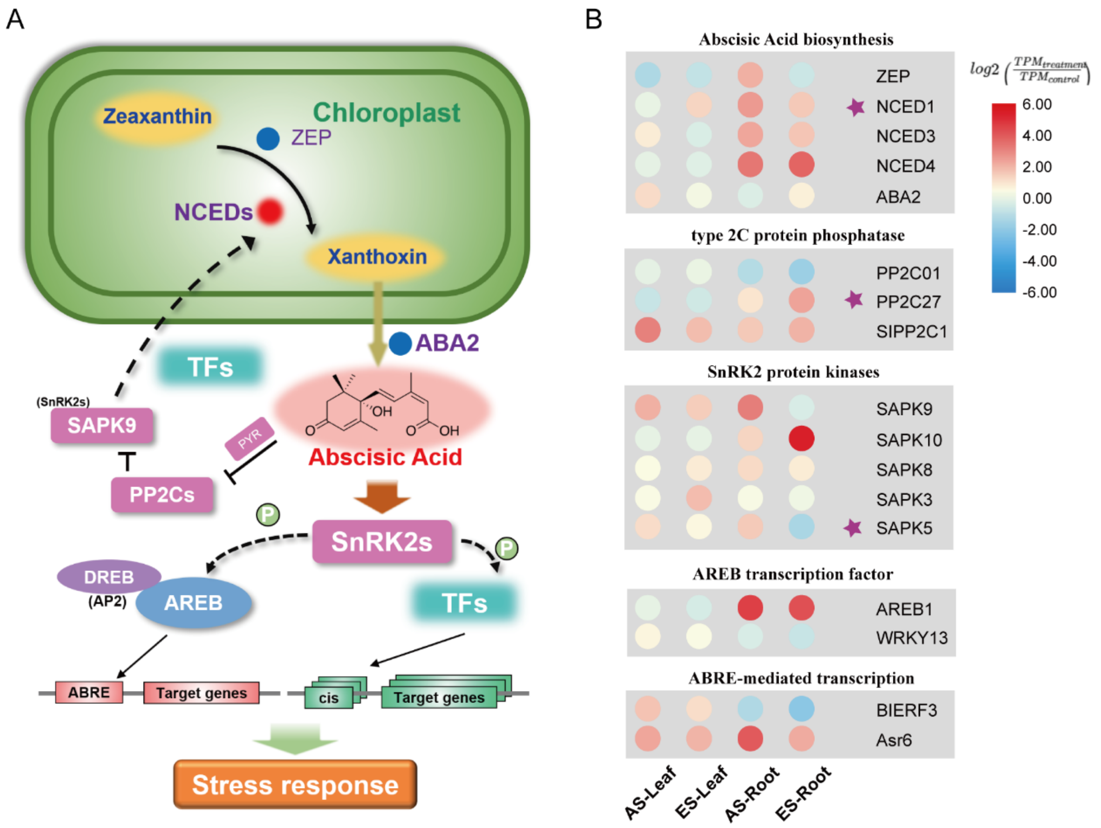


**Figure S24.** Different expression patterns of ABA signaling pathway genes in *Brachypodium stacei* leaf and root tissues of samples from the African Slope (AS) and European Slope (ES) populations at ECI under drought stress treatment. (A) Scheme of the ABA signaling pathway. (B) Heatmaps of differential expression patterns of genes of AS and ES leaf and root tissue samples for relevant genes of the ABA signaling pathway routes. Each cell represents the Log2 fold change of DEGs between drought (treatment) and well-watered (control) conditions ($TPM_{Treatment}/TPM_{Control}$); the color codes for these values are indicated in the corresponding chart. Genes located in selective regions are marked with a star.

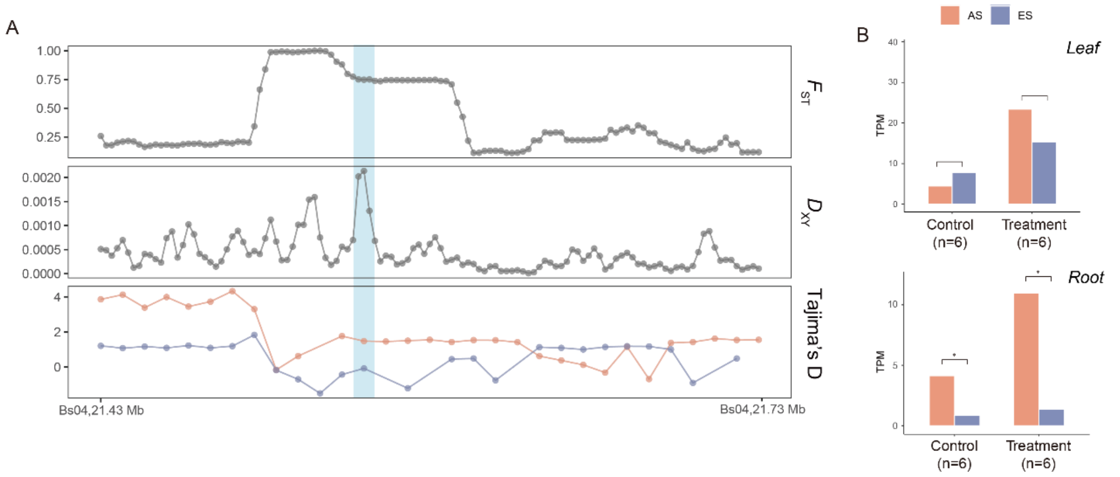


**Figure S25.** *Brachypodium stacei GL1-2* gene. A) Chromosomal location of a *Glossy1* (GL1) *Brachypodium stacei* orthologous *GL1-2* gene in chromosome Bs04 of the Bsta-ECI local reference genome. Pairwise $F_{ST}$, $D_{XY}$ and Tajima's D values from *GL1-2* sequences in the sequenced African Slope (AS) and European Slope (ES) individuals from Evolution Canyon I. (B) Expression levels of *GL1-2* in transcripts per million (TPM) in leaf and root samples of *B. stacei* samples from the AS and ES populations under well-watered (control) vs drought stress (treatment) conditions. The expression levels of *GL1-2* were significantly higher in AS than in ES root samples under drought and control conditions and in leaf samples under drought conditions, * p-value < 0.05 (Wilcoxon rank sum test).

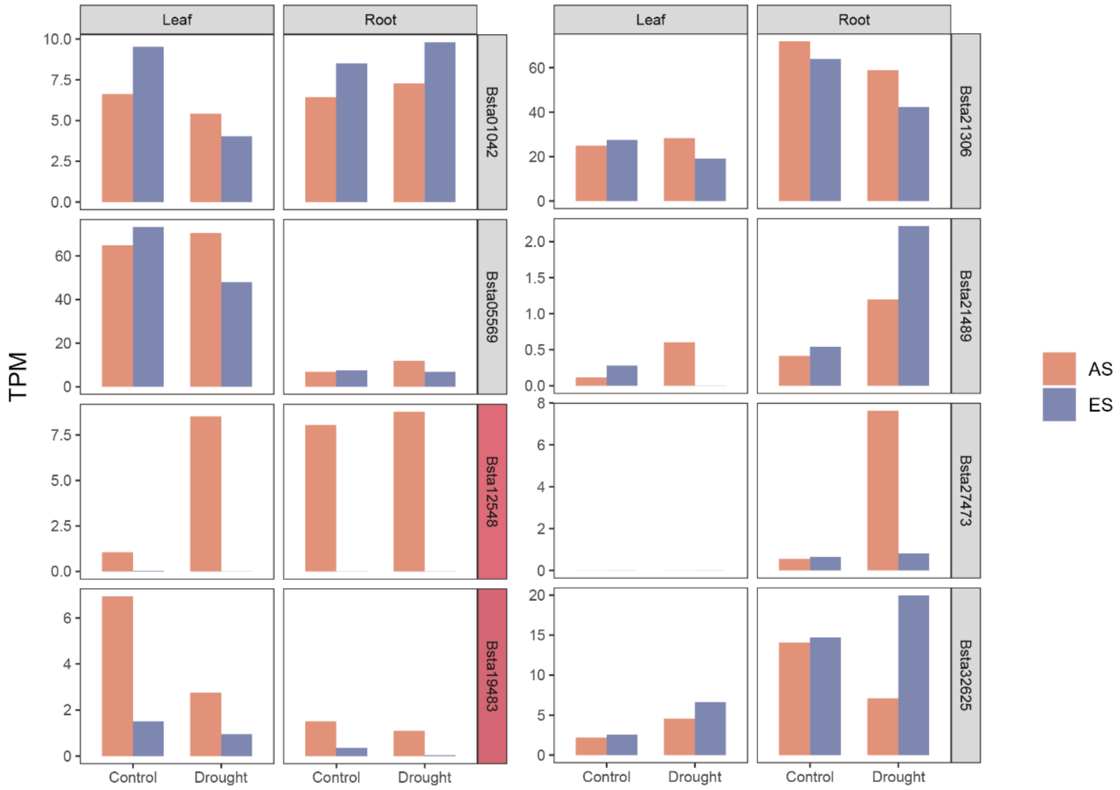


**Figure S26.** The expression level of 9 genes nearby TIPs in the different *B. stacei* AS and ES population samples' tissues and conditions. The genes identified as DEG in all inter-population comparisons were marked as red. AS and ES color codes are indicated in the chart.

#### Supplementary Tables: *Brachypodium stacei*

**Table S1.** Characteristics of the genome sequencing data for the *Brachypodium stacei* (Bsta-ECI) local reference genome from Evolution Canyon I, Israel (ECI). (See Dataset S1).

| Method | Library type | Reads number | Data size (Gb) | Mean read length (bp) | Read N50 (bp) |
|---|---|---|---|---|---|
| Illumina (clean data) | Paired | 115,050,912 | 17.24 | 150 | - |
| Pacbio Hifi | Single | 1,639,829 | 25.01 | 15253 | 15357 |
| Hi-C (clean data) | Paired | 1,011,713,862 | 151.75 | 150 | - |

**Table S2.** Comparison of genome assembly parameters of *Brachypodium stacei* Bsta-ECI and the *B. stacei* ABR114 reference genome generated by the Joint Genome Institute (JGI; Phytozome https://phytozome-next.jgi.doe.gov/; *B. stacei* ABR114 v. 1.1).

| **Genome feature** | ***B. stacei* (ECI)** | ***B. stacei* (ABR114 v.1.1)** |
|---|---|---|
| Total length of contigs (Mb) | 256.71 | - |
| Total length of assemblies (Mb) | 248.99 | 234.14 |
| Gap number | 30 | 3,132 |
| Number of contigs | 79 | 3,244 |
| Contig N50 (Mb) | 10.19 | 0.23 |
| Number of scaffolds | 10 | 10 |
| Scaffold N50 (Mb) | 24.13 | 23.06 |
| LAI | 10.46 | 2.06 |
| GC content (%) | 45.53 | 44.71 |
| Percentage of anchoring (%) | 96.99 | - |

**Table S3.** Summary of Hi-C mapping parameters of the *Brachypodium stacei* Bsta-ECI genome from Evolution Canyon I.

| | ***B. stacei* Bsta-ECI** | |
|---|---|---|
| | **Read number** | **Percentage (%)** |
| Total read pairs | 502,167,104 | 100 |
| Mapped read pairs | 425,121,335 | 84.66 |
| Uniquely mapped read pairs | 313,511,156 | 62.43 |
| Valid interaction read pairs | 258,389,405 | 51.45 |

**Table S4.** Chromosome level comparisons of the assembled *Brachypodium stacei* Bsta-ECI from Evolution Canyon I to the reference genome *B. stacei* ABR114 v 1.1 (Phytozome; https://phytozome-next.jgi.doe.gov/).

*B. stacei*

| | Bsta-ECI | | ABR114 v. 1.1 | |
|---|---|---|---|---|
| | Chromosome Length (bp) | Gap Number | Chromosome Length (bp) | Gap Number |
| Bs01 | 32,090,747 | 4 | 29,836,869 | 313 |
| Bs02 | 29,360,157 | 4 | 27,560,134 | 242 |
| Bs03 | 27,566,100 | 0 | 25,035,503 | 339 |
| Bs04 | 25,728,233 | 2 | 24,459,551 | 277 |
| Bs05 | 24,136,534 | 3 | 22,826,389 | 279 |
| Bs06 | 23,726,300 | 3 | 21,437,706 | 416 |
| Bs07 | 22,169,934 | 7 | 20,606,078 | 310 |
| Bs08 | 22,261,700 | 3 | 20,433,684 | 355 |
| Bs09 | 21,987,753 | 1 | 20,329,407 | 271 |
| Bs10 | 19,969,809 | 2 | 18,553,268 | 314 |

**Table S5.** Summary of genome assembly validation and statistics of the newly assembled *Brachypodium stacei* Bsta-ECI genome from Evolution Canyon I and the reference genome *B. stacei* ABR114 v.1.1

| | *B. stacei* Bsta-ECI | *B. stacei* ABR114 v1.1 |
|---|---|---|
| BUSCOs (%) | 98.6 | 98.4 |
| Complete and single-copy BUSCOs (%) | 97.3 | 97.1 |
| Complete and duplicated BUSCOs (%) | 1.3 | 1.3 |
| Fragmented BUSCOs (%) | 0.4 | 0.5 |
| Missing BUSCOs (%) | 1.0 | 1.1 |
| Mapping statistics | | |
| Fraction of Mapped Illumina Data (%) | 99.51 | 97.24 |
| Fraction of Properly read pairs Mapped (%) | 99.16 | 96.41 |
| Regions of Coverage > 0x (%) | 99.74 | 98.85 |
| Regions of Coverage ≥ 10x (%) | 99.63 | 98.82 |
| K-mer | | |
| Base call accuracy (QV) | 39.79 | 40.18 |
| Completeness | 99.10 | 96.16 |

**Table S6.** Prediction of repetitive elements in the assembled *Brachypodium stacei* Bsta-ECI genome from Evolution Canyon I.

| Type | Repeat Size (bp) | Percent of genome (%) |
|---|---|---|
| **TRF[1]** | 18,301,727 | 7.13 |
| **RepeatMasker** | 50,048,364 | 19.50 |
| **RepeatProteinMask** | 16,203,285 | 6.51 |
| ***De novo*** | 97,141,916 | 37.84 |
| **Total** | 103,467,454 | 40.31 |

1: TRF: Tandem Repeat Finder.

**Table S7.** Summary of repetitive elements in the newly assembled *B. stacei* BstaECI genome and in the *B. stacei* ABR114 v 1.1 reference genome

| Repetitive element type | | Length(bp) of genome | | Percentage (%) of Repeat type | | Percentage (%) of genome covered by repeat type | |
|---|---|---|---|---|---|---|---|
| | | Bsta- ECI | ABR114 | Bsta- ECI | ABR114 | Bsta- ECI | ABR114 |
| **SINE** | | 611,446 | 436,828 | 0.591 | 0.57 | 0.238 | 0.19 |
| **LINE** | | 7,583,328 | 6,785,199 | 7.329 | 8.86 | 2.954 | 2.90 |
| | L1 | 7,492,395 | 6,756,750 | 7.241 | 8.45 | 2.919 | 3.21 |
| | Other | 210,321 | 29,953 | 0.2 | 0.04 | 0.08 | 0.01 |
| **LTR** | | 46,478,386 | 35,045,130 | 44.921 | 45.77 | 18.106 | 14.97 |
| | Copia | 16,894,953 | 10,043,204 | 16.329 | 13.12 | 6.581 | 4.29 |
| | Gypsy | 29,297,286 | 23,934,981 | 28.315 | 31.26 | 11.413 | 10.22 |
| | Other | 4,543,886 | 1,496,957 | 4.392 | 1.96 | 1.77 | 0.64 |
| **DNA** | | 18,532,599 | 16,158,902 | 17.912 | 21.1 | 7.219 | 6.90 |
| | CMC-EnSpm | 3,537,181 | 2,786,276 | 3.419 | 3.64 | 1.378 | 1.19 |
| | hAT-Ac | 3,979,100 | 897,819 | 3.846 | 1.17 | 1.55 | 0.38 |
| | hAT-Tip100 | 2,780,739 | 571,236 | 2.688 | 0.75 | 1.083 | 0.24 |
| | MuDR | 1,020,941 | 3,313,096 | 0.987 | 4.33 | 0.398 | 1.42 |
| | PIF-Harbinger | 443,859 | 2,724,305 | 0.429 | 2.44 | 0.173 | 1.01 |
| | Other | 7,339,227 | 6,325,645 | 7.093 | 8.26 | 2.859 | 2.70 |
| **Transposable elements(TEs) total** | | 73,205,759 | 58,426,059 | 70.75 | 76.3 | 28.52 | 24.95 |
| **Satellites** | | 126,999 | 14,724 | 0.123 | 0.02 | 0.049 | 0.01 |
| **Simple_repeats** | | 19,846,896 | 5,427,411 | 19.182 | 7.09 | 7.731 | 2.32 |
| **Unclassified** | | 29,067,671 | 16,443,949 | 28.09 | 21.47 | 10.38 | 7.02 |
| **Total repeats** | | 103,467,454 | 76,575,937 | 100 | 100 | 40.31 | 32.70 |

**Table S8.** Gene predictions and statistics in the assembled *B. stacei* Bsta-ECI genome from Evolution Canyon I, compared to those of the reference genomes of *B. stacei* ABR114 v. 1.1, , and *B. distachyon* Bdis-Bd21 generated by the Joint Genome Institute (Phytozome, https://phytozome-next.jgi.doe.gov/), and of *Oryza sativa* subsp. *japonica*, generated by National Center for Biotechnology Information (NCBI_Assembly: GCF_001433935.1).

| **Species** | **Gene Predicted** | **Average Gene Length (bp)** | **Average CDS Length (bp)** | **Average Exons per Gene** | **Average Exon Length (bp)** | **Average Intron Length (bp)** | **Complete BUSCO (%)** |
|---|---|---|---|---|---|---|---|
| Bsta-ECI | 32,951 | 2528.92 | 1159.56 | 4.59 | 252.13 | 380.47 | 99.1 |
| Bsta-ABR114 | 29,898 | 3334.01 | 1197.6 | 4.76 | 251.26 | 393.98 | 99.2 |
| Bdis-Bd21 | 34,310 | 3373.23 | 1117.39 | 4.39 | 254.36 | 403.07 | 99.5 |
| *O. sativa japonica* | 37,869 | 2986.76 | 982.71 | 3.74 | 262.43 | 426.67 | 99.7 |

**Table S9.** Functional annotation of the predicted genes for *Brachypodium stacei* Bsta-ECI genome from Evolution Canyon I.

| | **Database** | ***B. stacei* Bsta-ECI** | |
|---|---|---|---|
| | | **Number** | **Percentages (%)** |
| Total | | 32,951 | 100 |
| | NR | 30,315 | 92.00 |
| | InterPro | 29,646 | 89.96 |
| Annotated | Swissport | 20,785 | 63.07 |
| | TrEMBL | 30,895 | 93.76 |
| | EggNOG | 27,760 | 84.24 |
| Unannotated | | 872 | 2.64 |

**Table S10.** Transcription factor (TF) prediction in *Brachypodium stacei* Bsta-ECI genome from Evolution Canyon I.

| TF family | *B. stacei* **Bsta-ECI** |
|---|---|
| Alfin-like | 9 |
| AP2/ERF-AP2 | 25 |
| AP2/ERF-ERF | 124 |
| AP2/ERF-RAV | 4 |
| B3 | 52 |
| B3-ARF | 26 |
| BBR-BPC | 3 |
| BES1 | 7 |
| bHLH | 127 |
| bZIP | 85 |
| C2C2-CO-like | 9 |
| C2C2-Dof | 29 |
| C2C2-GATA | 29 |
| C2C2-LSD | 6 |
| C2C2-YABBY | 8 |
| C2H2 | 113 |
| C3H | 56 |
| CAMTA | 7 |
| CPP | 9 |
| CSD | 4 |
| DBB | 4 |
| DBP | 6 |
| DDT | 6 |
| E2F-DP | 10 |
| EIL | 6 |
| FAR1 | 88 |
| GARP-ARR-B | 8 |
| GARP-G2-like | 49 |
| GeBP | 15 |
| GRAS | 58 |
| GRF | 12 |
| HB-BELL | 14 |
| HB-HD-ZIP | 38 |
| HB-KNOX | 11 |
| HB-other | 12 |
| HB-PHD | 3 |
| HB-WOX | 13 |

| | |
|---|---|
| HRT | 1 |
| HSF | 25 |
| LFY | 1 |
| LIM | 6 |
| LOB | 26 |
| MADS-MIKC | 34 |
| MADS-M-type | 43 |
| MYB | 122 |
| MYB-related | 65 |
| NAC | 122 |
| NF-X1 | 2 |
| NF-YA | 7 |
| NF-YB | 13 |
| NF-YC | 13 |
| OFP | 30 |
| PLATZ | 15 |
| RWP-RK | 16 |
| S1Fa-like | 1 |
| SBP | 17 |
| SRS | 6 |
| STAT | 1 |
| TCP | 22 |
| Tify | 15 |
| Trihelix | 28 |
| TUB | 12 |
| ULT | 1 |
| VOZ | 2 |
| Whirly | 2 |
| WRKY | 84 |
| zf-HD | 21 |
| All | 1,838 |
| Percentage of total gene predictions | 5.5% |

**Table S11.** Summary of genomic variants obtained in the 41 *Brachypodium stacei* individuals studied in Evolution Canyon I.

| Chromosome | Length | SNP number | SNP rate(%) | INDEL number | INDEL rate (%) |
|---|---|---|---|---|---|
| Bs01 | 32,090,747 | 96,922 | 0.30 | 28375 | 0.09 |
| Bs02 | 29,360,157 | 65,230 | 0.22 | 20286 | 0.07 |
| Bs03 | 27,566,100 | 95,553 | 0.34 | 25950 | 0.09 |
| Bs04 | 25,728,233 | 74,013 | 0.29 | 23583 | 0.09 |
| Bs05 | 24,136,534 | 56,799 | 0.23 | 19288 | 0.08 |
| Bs06 | 23,726,300 | 63,596 | 0.27 | 19343 | 0.08 |
| Bs07 | 22,169,934 | 86,554 | 0.40 | 26363 | 0.12 |
| Bs08 | 22,261,700 | 77,838 | 0.35 | 23108 | 0.10 |
| Bs09 | 21,987,753 | 30,580 | 0.14 | 11920 | 0.05 |
| Bs10 | 19,969,809 | 75,266 | 0.38 | 22105 | 0.11 |
| All | 248,997,267 | 722,351 | 0.29 | 220321 | 0.09 |

**Table S12.** Summary of Structure Variants (SVs) detected among the 41 *Brachypodium stacei* resequenced genomes of Evolution Canyon I (Del: deletion; Dup: duplication; Inv: inversion; Ins: insertion; Tra: translocation; UNK: unknow) relative to the Bsta-ECI local reference genome.

| SVs Len | Del | Dup | Inv | Ins | Tra | UNK |
|---|---|---|---|---|---|---|
| 0-50bp | 0 | 0 | 2 | 0 | 5954 | 0 |
| 50-100bp | 1352 | 97 | 32 | 1014 | 0 | 0 |
| 100-1000bp | 2023 | 156 | 90 | 560 | 0 | 0 |
| 1000-10000bp | 1016 | 242 | 104 | 0 | 0 | 0 |
| 10000+bp | 283 | 232 | 654 | 0 | 0 | 0 |

**Table S13.** Comparison of performances of seven alternative models of inference of evolutionary demographic history of *Brachypodium stacei* AS and ES populations in Evolution Canyon I conducted with *∂a∂i* using forward simulation and residuals analysis. The best asym_mig_twoepoch model was selected based on likelihood and AIC (Akaike Information Criterion) values. Each model was run 100 rounds using the dadi_pipeline (see methods); best likelihood and AIC values for each model are shown.

| **models** | **Best likelihood** | **Best AIC** |
|---|---|---|
| asym_mig_twoepoch | -22639.63 | 45295.26 |
| no_mig | -34200.1 | 68406.2 |
| sym_mig | -25200.15 | 50408.3 |
| asmy_mig | -23854.14 | 47718.28 |
| sym_mig_twoepoch | -24925.77 | 49863.54 |
| ancestor_divergence | -26997.49 | 54004.98 |
| second_contact | -24060.63 | 48131.26 |

**Table S14.** Parameters of the best asym_mig_twoepoch evolutionary demographic history model for the *Brachypodium stacei* AS and ES populations of Evolution Canyon I estimated from *∂a∂i*. NA, real population size of MRCA; nu1 and nu2, real population sizes of AS and ES populations; m12a and m21a, migration rates from AS to ES and from ES to AS in time slice T1; m12b and m21b, migration rates from AS to ES and from ES to AS in time slice T2; T1, initial time of epoch T1 (to T2); T2, initial time of epoch T2 (to present).

| Parameter | ∂a∂i output | Transference to readable |
|---|---|---|
| NA | 1 | 13421.18 |
| nu1 | 0.3357 | 4505.49 |
| nu2 | 0.1978 | 2654.71 |
| m12a | 3.1526 | 0.00011 |
| m21a | 1.7528 | 6.530 $e^{-5}$ |
| m12b | 1.542 | 5.744 $e^{-5}$ |
| m21b | 1.3721 | 5.111 $e^{-5}$ |
| T1 | 0.1131 | 10167.89 |
| T2 | 0.3788 | 3035.87 |

**Table S15.** Distribution of *Brachypodium stacei* AS and ES resequenced genomes 'genomic island' based on the top 5% of the $F_{ST}$ and $D_{XY}$ values obtained through linked sweeping located on each chromosome.

| Chr | Island windows distributed over one chromosome | windows distributed over one chromosome | all island windows | all windows | P-value (chisq.test) | Island length >70 kb (long) | Island length 40-70 kb (middle) | Island length 10-30 kb (short) |
|---|---|---|---|---|---|---|---|---|
| Bs01 | 41 | 2,977 | 276 | 22,932 | 0.476 | 1 | 3 | 17 |
| Bs02 | 28 | 2,648 | 276 | 22,932 | 0.578 | 0 | 2 | 14 |
| Bs03 | 10 | 2,603 | 276 | 22,932 | 2.77E-04 | 0 | 1 | 6 |
| Bs04 | 25 | 2,388 | 276 | 22,932 | 0.571 | 0 | 1 | 14 |
| Bs05 | 34 | 2,169 | 276 | 22,932 | 0.178 | 1 | 3 | 8 |
| Bs06 | 52 | 2,214 | 276 | 22,932 | 1.31E-05 | 1 | 4 | 13 |
| Bs07 | 31 | 2,118 | 276 | 22,932 | 0.355 | 2 | 1 | 6 |
| Bs08 | 18 | 2,146 | 276 | 22,932 | 0.167 | 0 | 1 | 12 |
| Bs09 | 33 | 1,770 | 276 | 22,932 | 0.023 | 0 | 3 | 10 |
| Bs10 | 4 | 1,899 | 276 | 22,932 | 1.47E-04 | 0 | 1 | 0 |

**Table S16.** Detection of a putative *Brachypodium stacei* ortholog to rice *OsNCED1,* a key regulatory gene in ABA biosynthesis, through reciprocal Blast searches. Only the top 5 hits of each search are shown. The best match corresponds to *Bsta13013* (84.3% identity).

| query id | subject id | Percentage of identity | evalue | bitscore |
|---|---|---|---|---|
| Os02g0704000 | Bsta13013 | 84.3 | 1.30E-270 | 929.5 |
| Os02g0704000 | Bsta07047 | 42.3 | 5.40E-104 | 375.9 |
| Os02g0704000 | Bsta19139 | 40.6 | 1.50E-98 | 357.8 |
| Os02g0704000 | Bsta30194 | 39.5 | 2.30E-94 | 344 |
| Os02g0704000 | Bsta30080 | 36.5 | 9.30E-88 | 322 |
| Bsta13013 | Os02g0704000 | 84.3 | 0 | 918 |
| Bsta13013 | Os12g0435200 | 66.5 | 2.38E-255 | 714 |
| Bsta13013 | Os03g0645900 | 42.6 | 1.44E-117 | 363 |
| Bsta13013 | Os07g0154100 | 42.3 | 3.58E-114 | 353 |
| Bsta13013 | Os12g0640600 | 37.9 | 2.14E-94 | 300 |

**Table S17.** Location of 16 SNPs in the 5'-upstream regulatory region of the *Brachypodium stacei Bsta13013* gene**,** a putative ortholog of rice NCED1, separating the AS and ES population samples' sequences, that show different haplotypes between them. AS-ES allelic frequencies of polymorphic *Bsta13013* positions were compared relative to the *Bsta13013* sequence of the *B. stacei* Bsta-ECI local reference genome.

| Bsta-ECI chromosome | Bsta-ECI position | AS representative seq frequence | AS allelic frequence | ES representative seq frequence | ES allelic frequence |
|---|---|---|---|---|---|
| Bs04 | 5823937 | T:0.791 | G:0.208 | T:0 | G:1 |
| Bs04 | 5824072 | C:1 | T:0 | C:0 | T:1 |
| Bs04 | 5824375 | A:0.8 | T:0.2 | A:0 | T:1 |
| Bs04 | 5824376 | C:0.8 | G:0.2 | C:0 | G:1 |
| Bs04 | 5824706 | C:1 | T:0 | C:0 | T:1 |
| Bs04 | 5824961 | G:0.8 | C:0.2 | G:0 | C:1 |
| Bs04 | 5825020 | A:0.8 | G:0.2 | A:0 | G:1 |
| Bs04 | 5825029 | C:0.8 | A:0.2 | C:1 | A:0 |
| Bs04 | 5825092 | A:0.8 | G:0.2 | A:1 | G:0 |
| Bs04 | 5825111 | T:0.8 | C:0.2 | T:0 | C:1 |
| Bs04 | 5825184 | A:0.8 | G:0.2 | A:0 | G:1 |
| Bs04 | 5825199 | A:0.8 | T:0.2 | A:0 | T:1 |
| Bs04 | 5825222 | T:0.8 | C:0.2 | T:0 | C:1 |
| Bs04 | 5825237 | A:1 | T:0 | A:0 | T:1 |
| Bs04 | 5825415 | A:0.8 | G:0.2 | A:0 | G:1 |
| Bs04 | 5825601 | G:0.8 | A:0.2 | G:0 | A:1 |
